# Insecticide resistance in *Myzus persicae* collected from sweet pepper

**DOI:** 10.1101/2024.09.19.613823

**Authors:** Mariska M. Beekman, Xinyan Ruan, Marcel Dicke, Bas J. Zwaan, Bart A. Pannebakker, Eveline C. Verhulst

**Affiliations:** Laboratory of Genetics, Wageningen University & Research, Wageningen, the Netherlands; Laboratory of Entomology, Wageningen University & Research, Wageningen, the Netherlands

## Abstract

Populations of the green peach aphid, *Myzus persicae*, rapidly develop resistance to insecticides applied in agriculture, necessitating regular resistance monitoring of pest populations. Previous research identified two dominant multilocus genotypes (MLGs) in conventional (using insecticides + biological control agents) Dutch sweet pepper greenhouses: MLG-A under pymetrozine application and MLG-R under flonicamid application. This suggests positive selection for the genotypes by the application of these insecticides. However, no resistance of *M. persicae* to these insecticides has been reported yet. To investigate whether the insecticides were selectively driving the emergence of these specific MLGs, we compared the sensitivity of MLG-A and MLG-R to pymetrozine and flonicamid to that of six other genotypes from the same crop system. Additionally, we screened the *M. persicae* populations from Dutch sweet pepper greenhouses for known mutations conferring resistance to carbamates, pyrethroids, neonicotinoids, and tetronic and tetramic acid derivatives. Our results show that both MLG-A and MLG-R are less sensitive to pymetrozine compared to the other genotypes investigated. Furthermore, full mortality for MLG-R, the genotype least sensitive to flonicamid, was not achieved at the recommended field dose for this insecticide. Resistance mutations for carbamates and pyrethroids were prevalent among the MLGs, including MLG-A and MLG-R, while mutation A2226V, which is linked to resistance to tetronic and tetramic acid derivatives, was absent. Notably, the neonicotinoid resistance mutation R81T was found only in MLG-R, making this the northernmost detection of R81T in *M. persicae* to date. This study shows that various resistance mechanisms can accumulate in a single aphid genotype and that insecticides likely play a role in selecting for the dominant genotypes of *M. persicae* in conventional greenhouses.

## Introduction

The green peach aphid, *Myzus persicae* (Sulzer), ranks among the most notorious agricultural pests worldwide. It is characterised by a cosmopolitan distribution (Margaritopoulos et al., 2009) and broad host range (Blackman and Eastop, 2000), and it can transmit many plant viruses (Nault, 1997; van Emden and Harrington, 2017). Furthermore, it exhibits a remarkable ability to quickly develop resistance to insecticides (Bass and Nauen, 2023; Bass et al., 2014), with reported resistance to 87 different compounds as of June 2024, covering almost all available insecticidal groups with aphicidal activity (Mota-Sanchez and Wise, 2024). Notably, *M. persicae* mostly reproduces through cyclical parthenogenesis, normally reproducing sexually once a year on its primary host, *Prunus* spp., where the eggs overwinter (Blackman, 1974). However, continuously asexually reproducing lineages also exist, either in the absence of environmental cues that trigger sexual reproduction, or as obligate asexuality. Because of its predominantly parthenogenetic life cycle, cosmopolitan distribution and its broad host range, *M. persicae* maintains an enormous effective population size, and with it, a high mutation supply. When a new beneficial mutation arises, parthenogenetic reproduction facilitates the rapid increase in the numbers of clone-mates (individuals all originating from a single parthenogenetic ancestor) carrying this mutation. When these clones skip sexual reproduction, enormous populations of clone-mates can form that can span large geographical areas and persist over long periods of time (Fenton et al., 2005; Fenton et al., 2010; Vorburger et al., 2003).

Although biological control (biocontrol) strategies are increasingly used to sustainably manage aphid populations, chemical insecticides are still commonly applied for controlling *M. persicae* in agricultural systems worldwide. A variety of insecticide classes with different modes of action (MoA), as classified by the Insecticide Resistance Action Committee (IRAC), are available for controlling aphids. However, the rapid emergence of insecticide resistance in this pest presents a significant challenge, and to date, at least seven independent biochemical and molecular resistance mechanisms have been described for *M. persicae* (reviewed by Bass et al., 2014). These include the enhanced expression of detoxifying esterases (Devonshire and Moores, 1982; Field et al., 1988), elevated levels of the detoxifying enzyme cytochrome P450 (Puinean et al., 2010), and various mutations in the genes encoding the target sites of different insecticidal groups. Together, these mechanisms lead to resistance to compounds belonging to the organophosphates (Devonshire and Moores, 1982; Field et al., 1988), cyclodienes (Anthony et al., 1998), carbamates (Andrews et al., 2004; Devonshire and Moores, 1982; Field et al., 1988; Nabeshima et al., 2003), pyrethroids (Eleftherianos et al., 2008; Fontaine et al., 2011; Panini et al., 2014), neonicotinoids (Bass et al., 2011) and tetronic and tetramic acid derivatives, also known as cyclic ketoenols (Singh et al., 2021; Umina et al., 2022).

Two insecticides noteworthy for their lack of reported resistance in *M. persicae* are pymetrozine and flonicamid (Mota-Sanchez and Wise, 2024). Moreover, both specifically target hemipterans, making them compatible with non-hemipteran biocontrol agents (Jansen et al., 2011; but see Fytrou et al., 2017; Joseph et al., 2011). This compatibility is crucial in crop systems that integrate biocontrol with chemical insecticides, as insecticides can have many negative effects of beneficial insect communities (Cloyd, 2012; Desneux et al., 2007). One cropping system where biocontrol and chemical insecticides are frequently integrated, and in which *M. persicae* is very problematic, is sweet pepper production in the Netherlands (Beekman et al., 2022; Messelink et al., 2020). Pymetrozine has been extensively utilised to control *M. persicae* in Dutch sweet pepper crops up until its EU-wide ban at the end of 2019 due to the potential endocrine disrupting properties and pollution of groundwater by its metabolites (European Commission, 2018). Despite the ban within the EU, pymetrozine continues to be widely used elsewhere (Pesticide Action Network International, 2022). Since 2020, flonicamid replaced pymetrozine as the primary insecticide for controlling *M. persicae* in Dutch sweet pepper greenhouses (Ctgb, 2024).

Alarmingly, reports from autumn 2021 and spring 2022 indicated significant difficulties in controlling *M. persicae* (Biobest Group, 2022; Glastuinbouw Nederland, 2022). Concurrently, our previous study on the population genetic structure of *M. persicae* in Dutch sweet pepper revealed the dominance of a specific *M. persicae* genotype in conventional (biocontrol + insecticides) greenhouses, which was absent from organic (biocontrol only) greenhouses (Beekman et al., 2024). This same study showed that in 2019 a different *M. persicae* genotype dominated in conventional greenhouses when pymetrozine was used, which was only moderately prevalent in organic greenhouses. The prevalence of dominant genotypes of *M. persicae*, alongside reported challenges in chemically controlling this aphid species, begs the question whether the dominating genotypes are less sensitive to the applied insecticides compared to other genotypes, even though resistance to neither pymetrozine nor flonicamid has been reported for *M. persicae* (Mota-Sanchez and Wise, 2024).

For effective *M. persicae* control in crop systems that integrate biocontrol with chemical insecticides, it is essential to balance biocontrol agents, insecticides, and resistance mitigation strategies. Compatibility of biocontrol agents and insecticides is generally based on the lethal effects of the insecticides on biocontrol agents. However, the numerous sublethal effects that insecticides have on beneficial arthropods have become increasingly clear (Cloyd, 2012; Desneux et al., 2007). Ideally, selective insecticides, which are implied to cause minimal harm to beneficial insects, are used (Roubos et al., 2014). However, the limited availability of such insecticides makes it challenging to rotate insecticides with different MoA, a core principle of resistance mitigation promoted by IRAC (https://www.irac-online.org). Resistance development of pests to selective insecticides is particularly problematic, as it might lead growers to apply more broad-spectrum insecticides, which can negatively affect biocontrol agent communities. It is thus crucial to regularly monitor the susceptibility of pest populations of *M. persicae* to the remaining effective insecticides, especially to selective ones.

In this study we evaluated the susceptibility of the *M. persicae* population from Dutch sweet pepper greenhouses to various insecticides, both at the genetic and phenotypic levels. Specifically, we: 1) tested whether the two genotypes that dominated in conventional Dutch sweet pepper greenhouses in 2019 and 2022 display reduced sensitivity to pymetrozine and flonicamid respectively, compared to other *M. persicae* genotypes from the same crop system, and 2) assessed the prevalence of known mutations involved in resistance to carbamates, pyrethroids, neonicotinoids, and tetronic and tetramic acid derivatives within this population. Lastly, we discuss the implications of our results for the selection of dominant genotypes in Dutch sweet pepper greenhouses.

## Materials and methods

### Pymetrozine and flonicamid sensitivity assays

#### Aphid lines

Laboratory aphid lines (Table 1) were started from single parthenogenetic females and cultured in sterile polypropylene culture vessels (Lab Associates, Oudenbosch, the Netherlands), 80 mm in height and having a bottom diameter of 92 mm, closed with nylon-screened Donut Lids (BDC0001-1; Bugdorm, MegaView Science, Taichung, Taiwan). Aphids were reared on sweet pepper (*Capsicum annuum* L.) leaf discs embedded in 1% agarose, stored upside down and maintained at 15 °C, under 16:8 light:dark photoperiod, and at 60% relative humidity in an incubator (Sanyo MLR-352H, PHC Europe, Etten-Leur, the Netherlands). The aphid lines were genotyped with the use of five microsatellite markers, as described in Beekman et al. (2024).

**Table 1.** Collection date and location of the living aphid lines used in the insecticide sensitivity assays. Aphid ID refers to the aphid line names. MLG represents the multilocus genotype of the aphid line, as determined in Beekman et al. (2024). Every greenhouse identifier represents a unique greenhouse, with those starting with ‘C’ being conventional and those starting with ‘O’ being organic. Greenhouse identifiers details can be found in Supplementary Table S1 of Beekman et al. (2024). Aphid lines are used in pymetrozine and flonicamid experiment 1 (PF1) and/or flonicamid experiment 2 (F2).

| Aphid ID | Collection date | Greenhouse identifier | MLG | In experiment |
| --- | --- | --- | --- | --- |
| A(C9-2019) | 19-Jul-19 | GH-C9 | A | PF1 |
| A(C7-2021) | 07-Jul-21 | GH-C7 | A | PF1, F2 |
| R(C13-2021) | 01-Oct-21 | GH-C13 | R | PF1, F2 |
| R(C14-2022) | 29-Mar-22 | GH-C14 | R | F2 |
| R(C15-2022) | 29-Mar-22 | GH-C15 | R | F2 |
| R(C16-2022) | 14-Apr-22 | GH-C16 | R | F2 |
| J(O7-2019) | 13-Jun-19 | GH-O7 | J | PF1 |
| P(O9-2019) | 18-Jun-19 | GH-O9 | P | PF1, F2 |
| K(C2-2021) | 14-May-21 | GH-C2 | K | PF1 |
| V(C2-2021) | 14-May-21 | GH-C2 | V | PF1 |
| T(C12-2021) | 28-May-21 | GH-C12 | T | PF1 |
| T(C12-2021)* | 16-Jun-21 | GH-C8 | T | PF1 |
| B(C4-2021) | 09-Jun-21 | GH-C4 | B | PF1 |

\* Note: The asterisk (\*) denotes an aphid line initially genotyped as MLG-A. However, after the experiments were conducted, genotyping to double-check the MLGs showed these aphids had MLG-T. We assume that this line was accidentally swapped with aphids from line T(C12-2021) at some point during the aphid rearing before the start of the experiments. Therefore, throughout this study, line T(C12-2021)\* is treated as MLG-T.

Prior to the start of the insecticide sensitivity assays, the population sizes of all aphid lines were boosted by transferring them to whole sweet pepper plants in nylon mesh rearing cages (4F3074; Bugdorm, MegaView Science, Taichung, Taiwan), maintained at 19 °C, under 16:8 light:dark photoperiod, and at 60% relative humidity. Regular monitoring of the multilocus genotypes (MLGs) of the aphid populations was performed using microsatellites as described in Beekman et al. (2024) to prevent accidental mixing of populations.

#### Pymetrozine and flonicamid experiment 1 (PF1)

The goal of this experiment was to compare the sensitivity of the various *M. persicae* MLGs (Table 1) to both pymetrozine and flonicamid, specifically investigating whether MLG-A and MLG-R, which dominated Dutch sweet pepper greenhouses in 2019 and 2022, respectively, are less sensitive to these insecticides compared to other *M. persicae* MLGs from the same crop system.

Leaf-dip bioassays were conducted following a modified version of the Insecticide Resistance Action Committee (IRAC) protocol 019 v. 3.4 (available at https://irac-online.org/methods/). These assays assessed the sensitivity of 10 *M. persicae* lines (eight MLGs), measured as mortality rates, to the insecticides pymetrozine and flonicamid. In short, for each line, adult aphids were exposed during 24 h to sweet pepper leaf discs dipped in various concentrations of the insecticides. Subsequently, the survival of the nymphs born within this time frame was assessed at 96 h after initial placement of the adults. Each treatment was replicated three times, with one replicate per temporal block.

Leaf-dip bioassays were conducted using the active ingredients pymetrozine and flonicamid, rather than commercial products, necessitating the addition of a solvent for pymetrozine, and a wetting agent for both insecticides, to ensure the water-based insecticide solutions spread evenly over the plant leaves. Pymetrozine (46119; Sigma-Aldrich, Zwijndrecht, the Netherlands) was prepared at 10 mg/mL in dimethylsulfoxide (DMSO) via sonication. The stock solution was used freshly for the first replicate and stored at −20 °C until use for the subsequent replicates. Flonicamid (32509; Sigma-Aldrich) was prepared at 1.5 mg/mL in ultrapure water with fresh stock solutions prepared for each experimental replicate. Various testing concentrations were derived from stock solutions in ultrapure water, with the addition of 0.01% Silwet® L-77 (Phytotechnology Laboratories, Kansas, USA) as a wetting agent. Based on earlier studies, the test concentrations were 2, 5, 10, 20, and 40 mg/L for pymetrozine (Foster et al., 2002; Margaritopoulos et al., 2010) and 2, 4, 8, 16, and 32 mg/L for flonicamid (Cho et al., 2011; Morita et al., 2007). For the pymetrozine treatments, additional DMSO was added to maintain a consistent final concentration of 0.4%. Control treatments consisted of ultrapure water with 0.01% Silwet® L-77 for flonicamid and an extra 0.4% DMSO for the pymetrozine control treatment. Sweet pepper leaf discs (90 mm diameter) were dipped into the various solutions for 10 s per side and left to air dry. Subsequently, the leaves were placed abaxial side up in 100 x 40 mm polystyrene insect breeding dishes (A19642; Novolab, Geraardsbergen, Belgium) on a 1% agarose layer. On each bioassay, 15-20 adult aphids were placed and left for 24 h at 18 °C, 16:8 light: dark photoperiod, and 60% humidity to produce offspring. After this period, the adults were removed and offspring survival was assessed 96 h after introduction of the adults.

#### Flonicamid experiment 2 (F2)

Next, we aimed to assess the sensitivity of different aphid lines with MLG-R, collected from various greenhouses, to flonicamid, to make sure that sensitivity is equal across various lines with the same MLG. Survival rates of four aphid lines with MLG-R were compared with two lines with other MLGs (Table 1). This experiment followed a similar setup as PF1, with adjustments made to better replicate commercial greenhouse conditions, including flonicamid concentration, temperature, and exposure duration. Fresh stock solutions of flonicamid were prepared at 1.75 - 1.86 mg/mL for each replicate. Testing concentrations were 32 and 50 mg/L, with the former corresponding to the highest tested dose in PF1 to allow for comparisons, and the latter corresponding to the concentration of active ingredient (Teppeki; 10 g/100 L with 50% active ingredient flonicamid) as recommended for sweet pepper greenhouses by the Dutch Board for the Authorisation of Plant Protection Products and Biocides (Ctgb). A control treatment was included similar to the procedure in PF1. Bioassays were initiated with 15 adults, were kept at 20 °C, and offspring survival was assessed 168 h after introduction of the adults.

#### Genotype checking

To confirm the genotypes of the tested aphids we used the methods described in Beekman et al. (2024) to genotype a random subsample of the aphids from the control treatments (three for each replicate of experiment PF1 and four for each replicate of experiment F2). We assessed the genotype on the relevant subset of microsatellite markers by examining the amplified microsatellites on a 4% agarose gel (for more details and results see Suppl. File S1).

#### Statistical analyses

Statistical analyses were carried out using R v. 4.2.1 (R Core Team, 2022) with RStudio v. 2022.07.0 (RStudio Team, 2022). Results were considered reliable when, after 96 h for experiments PF1, and 168 h for experiment F2, mortality in the respective control treatment was below 15% (Z. Li et al., 2023). Mortality was calculated by dividing the number of dead aphids by the total number of observed aphids.

Because not all aphid lines reached 100% mortality after exposure to the highest concentrations of the insecticides, reliable LC50 values could not be computed. As an alternative, we tested for an association between the aphid lines and their sensitivity to the insecticides, expressed as mortality, with generalised linear models (GLMs). We used the *glm* function of the *lme4* package v. 1.1-34 (Bates et al., 2015) based on a binomial distribution with a logit-link to model aphid mortality. To quantify aphid mortality, we employed a *cbind* procedure, combining counts of dead and surviving aphids. To be able to compare the morality between aphid lines, both insecticide concentration and aphid line were included as fixed factors, as categorical variables. Including replicate as a random factor in the generalised linear mixed model (GLMM) using the *glmer* function did not improve the models, as determined by the Akaike Information Criterion (AIC) (Akaike, 1998) which showed less than a 2-point difference between the models (Burnham and Anderson, 2004). Consequently, this factor was omitted from the final models.

The potential impact of the variable number of nymphs subjected to each treatment on the mortality rates was tested by including the log of the total number of nymphs tested in a treatment as an offset term to the models. However, the higher AIC values of the models including the offset suggest that the total number of nymphs included in each treatment did not improve the models and thus did not additionally affect aphid mortality. To identify which aphid lines exhibited significant differences in their survival response to the insecticides, we used the *Anova* function from the *car* package v. 3.1-2 (Fox and Weisberg, 2018), followed by a *post hoc* Tukey’s test using the *multcomp* package v. 1.4-25 (Hothorn et al., 2008). Effects plots were made using the estimated effects of the response variables (aphid line and insecticide concentration) on aphid mortality, as based on the GLMs and computed with the *allEffects* function of the R package *effects* v. 4.2-2 (Fox, 2003). Graphs were computed with the *ggplot2* package v. 3.5.1 (Wickham, 2016), and adjusted using Inkscape v. 1.2.2 (Inkscape Project, 2022).

### Screening for insecticide resistance-inducing mutations

We screened the *M. persicae* population from Dutch sweet pepper greenhouses, collected between 2019 and 2022 (Beekman et al., 2024), for known insecticide resistance-inducing mutations. Specifically, we examined the following mutations: A2226V linked to resistance to tetronic and tetramic acid derivatives (Singh et al., 2021; Umina et al., 2022), mutation S431F conferring resistance to dimethyl-carbamates (Andrews et al., 2004; Benting and Nauen, 2004; Nabeshima et al., 2003), mutation R81T associated with resistance to neonicotinoids (Bass et al., 2011), and mutations in the *para*-orthologous sodium channel gene (henceforth called ‘*para*’) causing resistance to pyrethroids (Eleftherianos et al., 2008; Fontaine et al., 2011; Panini et al., 2014; Rinkevich et al., 2013) (Table 2).

**Table 2.** Mutations causing resistance to insecticides that are studied at the genetic level in this study. ‘Chemical’ describes the chemical (sub-)group the insecticides belong to, together with the group these insecticides are assigned to according to The Insecticide Resistance Action Committee (IRAC, https://irac-online.org). ‘Mutation(s)’ are the names of the mutations, according to the standard nomenclature for the description of substitutions in proteins. ‘Mutation in’ describes which protein is affected by the mutation. ‘Products’ describes the names of the commercial products (with active ingredient) belonging to this chemical group that are allowed in Dutch sweet pepper greenhouses in 2024 according to the Netherlands Board of the Authorisation of Plant Protection Products and Biocides (2024).

| Chemical (IRAC) | Mutation(s) | Mutation in | Products |
| --- | --- | --- | --- |
| Dimethyl carbamates (1A) | S431F | Acetylcholinesterase enzyme (AChE) | Pirimor (pirimicarb) |
| Pyrethroids (3A) | M918T, M918L, M918I, L1014F, L932F, F979S | Voltage-gated sodium channel (VGSC) (also known as *para*) | Spiro (pyrethrins), Decis<sup>1</sup> (deltamethrin) |
| Neonicotinoids (4A) | R81T | Nicotinic acetylcholine receptor (nAChR) β1-subunit | Gazelle (acetamiprid) |
| Tetronic and tetramic acid derivatives (23) | A2226V | Acetyl-CoA carboxylase enzyme (ACCase) | Oberon<sup>2</sup> (spiromesifen) |

<sup>1</sup> applied against almost all herbivorous pests other than aphids (Van Iperen, 2024) <sup>2</sup> used against whiteflies and spider mites but has limited efficacy against aphids (Van Iperen, 2024).

#### Aphid genetic materials

Twenty-seven unique *M. persicae* MLGs, identified as part of a previous study on *M. persicae* from Dutch sweet pepper greenhouses (Beekman et al., 2024), were analysed for mutations known to be involved in insecticide resistance. DNA, isolated in the previous study using a Chelex and proteinase K-based protocol as described in Beekman et al. (2022), was available for 26 of these MLGs, with the exception of MLG-X. This previous study also resulted in the genome sequences of nine of the 27 MLGs, specifically MLG-A, B, D, J, K, P, R, T, and X, which are aligned to the reference genome of clone G006 v. 3 (available at https://bipaa.genouest.org/is/aphidbase/). The nine genomes were manually scanned for all aforementioned insecticide resistance-inducing mutations using the Integrative Genomics Viewer (IGV) v. 2.12.3 (Robinson et al., 2011). The exact locations of the mutations in the G006 v.3 reference genome are listed in Suppl. Table S1.

The resistance genotypes for the 18 MLGs for which no genome was available were analysed with PCR screening. For these, one DNA sample per MLG was analysed. We assumed one sample per MLG was sufficient because whole-genome sequencing of multiple samples of the same MLG previously showed that five microsatellite markers were enough to distinguish true clone-mates in the Dutch sweet pepper greenhouse system (Beekman et al., 2024). To confirm our assumption, we tested 13 samples, all with MLG-A, for the presence of S431F and for mutations in *para* and found identical results for all 13 samples (Suppl. Files S2 and S3).

#### Mutation S431F (carbamates)

The DNA samples of the 18 MLGs for which no genomic data was available were screened for mutation S431F via allele-specific PCR, following a modified version of the protocol by Fontaine et al. (2011). Primer Mace-R-Rev exclusively binds to the mutant allele, resulting in PCR amplification only when the mutation is present. PCR reactions were carried out in 10 µL-volumes containing 1x GoTaq® reaction buffer (Promega, Southampton, UK), 0.1 mM dNTPs, 0.2 µM forward primer AChE-F2 (see Table 3 for all primers), 0.2 µM reverse primer Mace-R-Rev, 0.25u GoTaq® DNA polymerase and 1 µL DNA. The PCR program was as follows: 3 min at 95 °C, followed by two cycles of 94 °C for 30s, 55 °C for 45s and 72 °C for 75s, followed by two cycles with an annealing temperature of 53 °C, two cycles with 51 °C, eight cycles with 49 °C, 21 cycles with 47 °C and a final extension at 72 °C for 5 minutes. PCR products were visualised by gel electrophoresis on a 1% agarose gel stained with ethidium bromide. The zygosity for this mutation was not assessed, except for the lines that were whole genome sequenced.

**Table 3.**
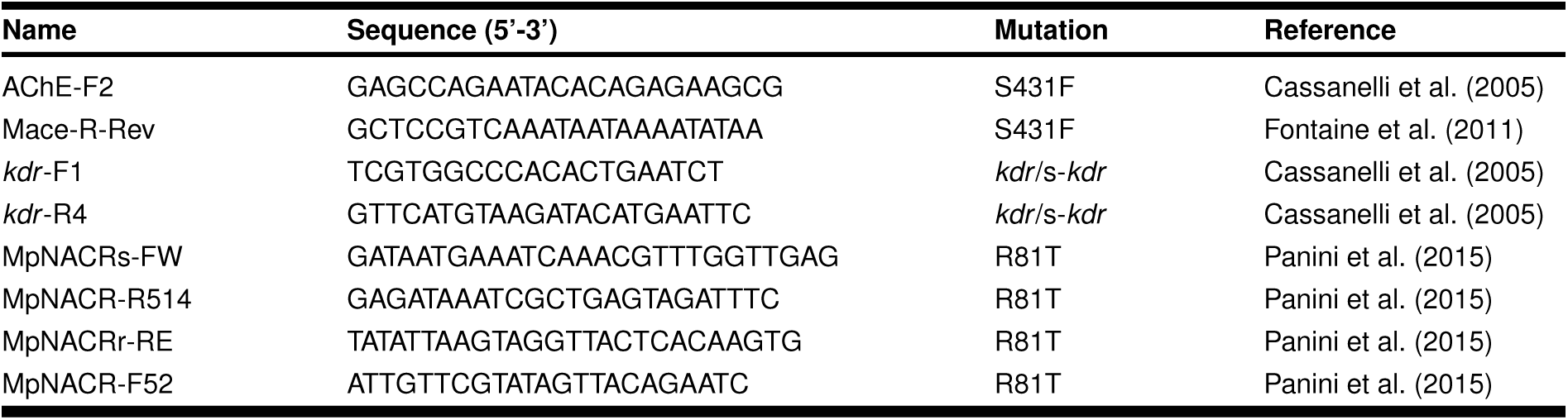
Primers used for insecticide resistance characterisation.

#### Mutations in para (pyrethroids)

We screened the 18 DNA samples for mutations in *para* by amplifying and sequencing a 578 bp fragment of the *para* gene encompassing domain II S4–S6, following a modified version of the protocol by Cassanelli et al. (2005). PCR reactions were carried out in 20 µL-volumes containing 1x GoTaq® reaction buffer (Promega), 0.1 mM dNTPs, 1 µM forward primer *kdr* -F1 (Table 3), 2 µM reverse primer *kdr* -R4 (Table 3), 0.5 u GoTaq® DNA polymerase and 2 µL DNA. The PCR program was as follows: 2 min at 94 °C, followed by 5 cycles of 94 °C for 40s, 60 °C for 45s and 72 °C for 1 min, followed by 10 cycles of touchdown PCR (annealing temperature −1 °C/cycle from 60 to 51 °C), followed by 25 cycles with an annealing temperature of 50 °C, and a final extension at 72 °C for 10 minutes. Successful PCR amplification was verified by gel electrophoresis. PCR products were purified using the NucleoSpin® Gel and PCR Clean-up kit (Macherey-Nagel, Dü ren, Germany) and send to Eurofins Genomics for Sanger sequencing of both strands. Electropherograms were analysed using Geneious Prime v. 2019.1.3 (BioMatters Ltd., Auckland, New Zealand).

#### Mutation R81T (neonicotinoids)

We attempted to detect the R81T substitution in the 18 DNA samples with allele-specific PCR by using the primers and protocol developed by Panini et al. (2014). This protocol combines four primers in a multiplex PCR. Specifically, primers MpNACR-FW and MpNACR-R514 (Table 3) are expected to amplify a 177 bp fragment of the WT allele, while MpNACRr-RE and MpNACR-F52 should amplify a 332 bp fragment of the mutant allele. Combined, primers MpNACR-R514 and MpNACR-F52 are anticipated to amplify a 508 bp fragment, common to both the WT and mutant alleles, serving as an internal control of the PCR. However, despite several optimisation attempts no clear PCR results were obtained (see Suppl. File S3 for attempted optimisation steps and results).

## Results

### Pymetrozine and flonicamid sensitivity assays

We aimed to investigating whether *M. persicae* lines with MLG-A and MLG-R (Table 1), which dominated Dutch sweet pepper greenhouses in 2019 and 2022, respectively, are less sensitive to pymetrozine and flonicamid, respectively, compared to other *M. persicae* MLGs from the same crop system.

#### Pymetrozine experiment 1

For aphid line A(C9-2019), control mortality was 17.4, 21.1 and 3.8% in the three replicates respectively (Suppl. Table S2). Given that control mortality exceeded the established 15% cut-off value (Z. Li et al., 2023) in two out of three replicates, this line was excluded from further analyses. Control mortality for all other lines ranged between 0 and 7.1% with an average of 0.9% (SD: ± 2.0) (Suppl. Table S2).

The number of nymphs included in the experiments, control treatments excluded, ranged from nine to 99, with an average of 34.7 (± 14.2) and there was significant variation in the number of nymphs tested between aphid lines, insecticide concentrations, and replicates (for details see Suppl. File S4).

Both the pymetrozine concentration (GLM: *χ*^2^(4, N = 135) = 360.03, *p <* 0.001; Figure 1B) and the aphid line (GLM: *χ*^2^(8, N = 135) = 447.78, *p <* 0.001; Figure 1C) had a significant effect on the mortality rate of *M. persicae* at four days post-treatment with pymetrozine. Mortality, as determined by GLM, increased with increasing concentration of pymetrozine, except at the two highest concentrations, where the mortality rates did not differ from each other (Figure 1B; for results of Tukey’s post hoc HSD tests, see Suppl. Table S3). Large differences in mortality rates between aphid lines were apparent, even at the lowest concentration of pymetrozine that we tested, because mortality at 2 mg/L pymetrozine, averaged over the three replicates, ranged from 6.1% (± 9.2) for line A(C7-2021) to 71.0% (± 6.6) for line T(C12-2021)* (Figure 1A; Suppl. Table S2). At the highest concentration of pymetrozine, 40 mg/L, this averaged mortality ranged from 60.6% (± 21.3) for line A(C7-2021) to 88.7% (± 6.8) for line J(O7-2019). The average mortality across all aphid lines ranged from 44.4% (± 22.3) at 2 mg/L to 76.4% (± 16.8) at 40 mg/L. Lines A(C7-2021) and R(C13-2021) exhibited the lowest sensitivity to pymetrozine with mortality rates, as determined by GLM, significantly lower than those of all other lines (Figure 1C; for results of Tukey’s post hoc tests, see Suppl. Table S3).

**Figure 1.**
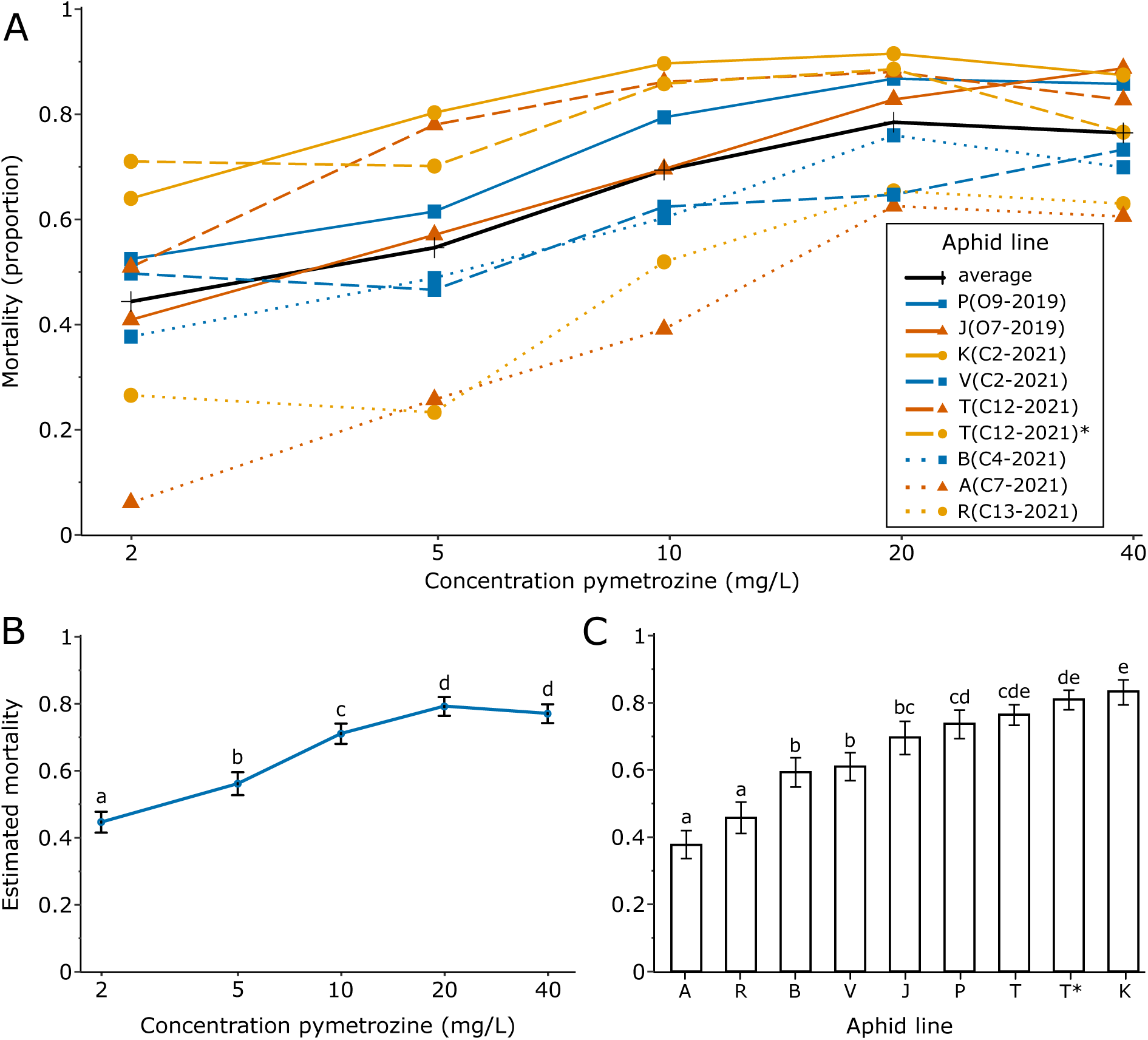
Responses of *Myzus persicae* to pymetrozine. A) Average measured mortality of nine different aphid lines (eight multilocus genotypes), over three replicates, when exposed for 96 h to five concentrations of pymetrozine (2, 5, 10, 20 and 40 mg/L). The black line represents the average mortality of all aphid lines together. B) Effects plot displaying the estimated effects of insecticide concentration and C) aphid line, ordered from lowest to highest average measured mortality, on aphid mortality, based on a generalised linear model. The x-axes of panels A and B are displayed on a logarithmic (base 2) scale. The error bars in panels B and C represent the standard error while the letters signify significant differences (*p <* 0.05) between treatments, as determined by Tukey’s post hoc testing. Aphid line IDs in panel C are abbreviated using the respective MLG of the line.

#### Flonicamid experiment 1

Mortality in the control treatments ranged from 0 to 5.6%, with an average of 0.6% (SD: ± 1.4), indicating that no line had to be excluded (Suppl. Table S4). The number of nymphs included in the experiments, control treatments excluded, ranged from 8 to 62, with an average of 27.1 (± 10.6), and differed significantly between replicates and aphid lines (for details see Suppl. File S4).

Both flonicamid concentration (GLM: *χ*^2^(4, N = 135) = 561.02, *p <* 0.001; Figure 2B) and aphid line (GLM: *χ*^2^(8, N = 135) = 299.99, *p <* 0.001; Figure 2C) had a significant effect on the mortality rate of *M. persicae* at four days post-treatment with flonicamid. Mortality increased with increasing concentration of flonicamid. Average mortality over all aphid lines ranged from 11.8% (± 9.1) at 2 mg/L to 67.3% (± 20.2) at 32 mg/L (Figure 2A; Suppl. Table S4). Lines R(C13-2021) and V(C2-2021) displayed the lowest mortality, while the lines with MLGs A, K, P, T displayed the highest mortality (Figure 2C; for results of Tukey’s post hoc tests, see Suppl. Table S5).

**Figure 2.**
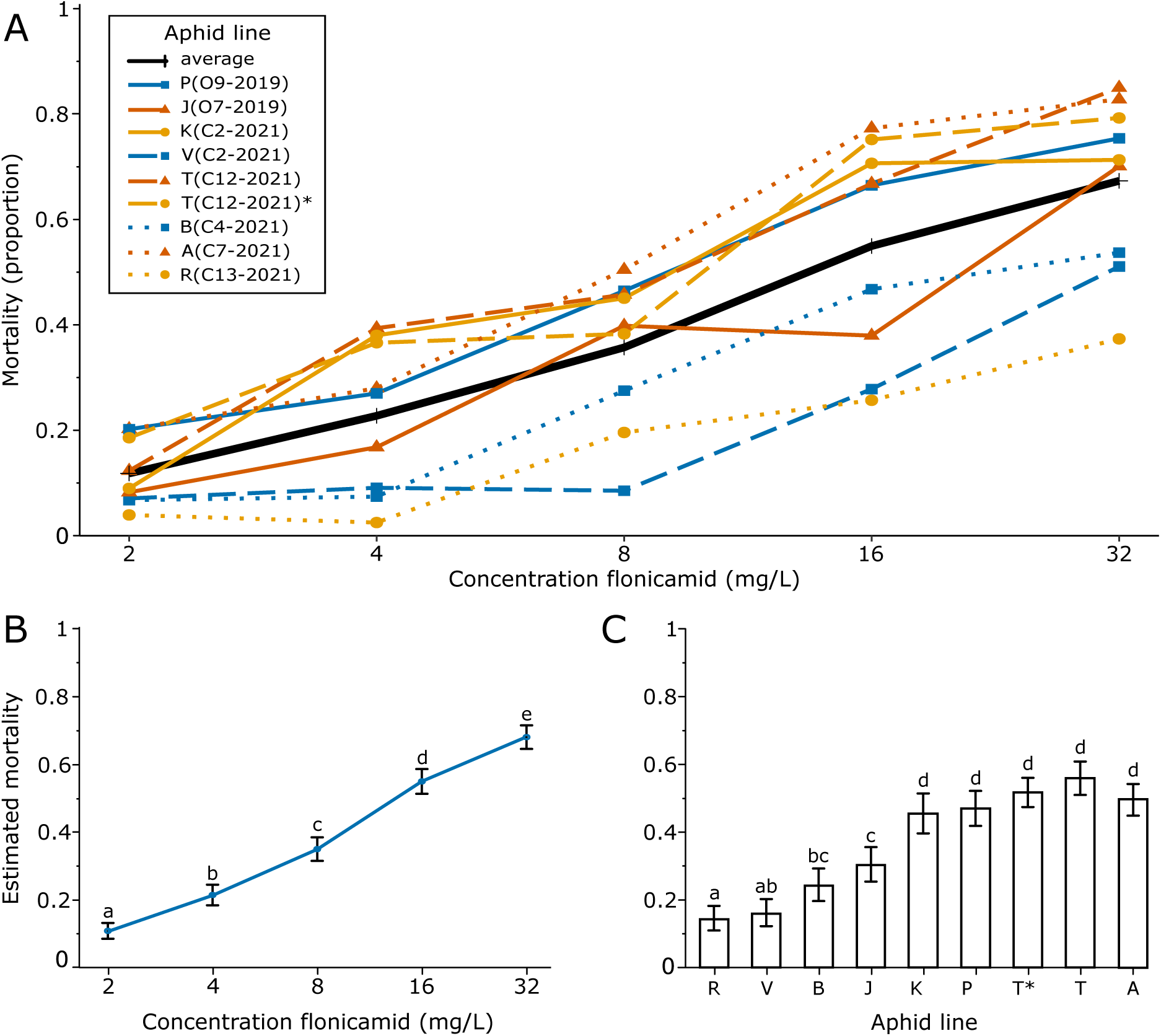
Responses of *Myzus persicae* to flonicamid. A) Average measured mortality of nine different aphid lines (eight multilocus genotypes) over three replicates, when exposed for 96 h to five concentrations of flonicamid (2, 4, 8, 16 and 32 mg/L). The black line represents the average mortality of all aphid lines together. B) Effects plot displaying the estimated effects of insecticide concentration and C) aphid line, ordered from lowest to highest average measured mortality, on aphid mortality, based on a generalised linear model. The x-axes of panels A and B are displayed on a logarithmic (base 2) scale. The error bars in panels B and C represent the standard error while the letters signify significant differences (*p <* 0.05) between treatments, as determined by Tukey’s post hoc testing. Aphid line IDs in panel C are abbreviated using the respective MLG of the line.

#### Flonicamid experiment 2

Next, we tested whether four lines with MLG-R from different greenhouses displayed similar levels of reduced sensitivity to flonicamid and we compared this with two non-MLG-R lines (Figure 3). Mortality in the control treatments ranged from 0 to 3.9% with an average of 0.6% (SD: ± 1.3) and thus no line had to be excluded (Suppl. Table S6). The number of nymphs tested in the treatments, control treatments excluded, ranged from 26 to 59 with an average of 42.8 (± 9.4) and was equal across insecticide concentrations, aphid lines and replicates (Suppl. File S4).

**Figure 3.**
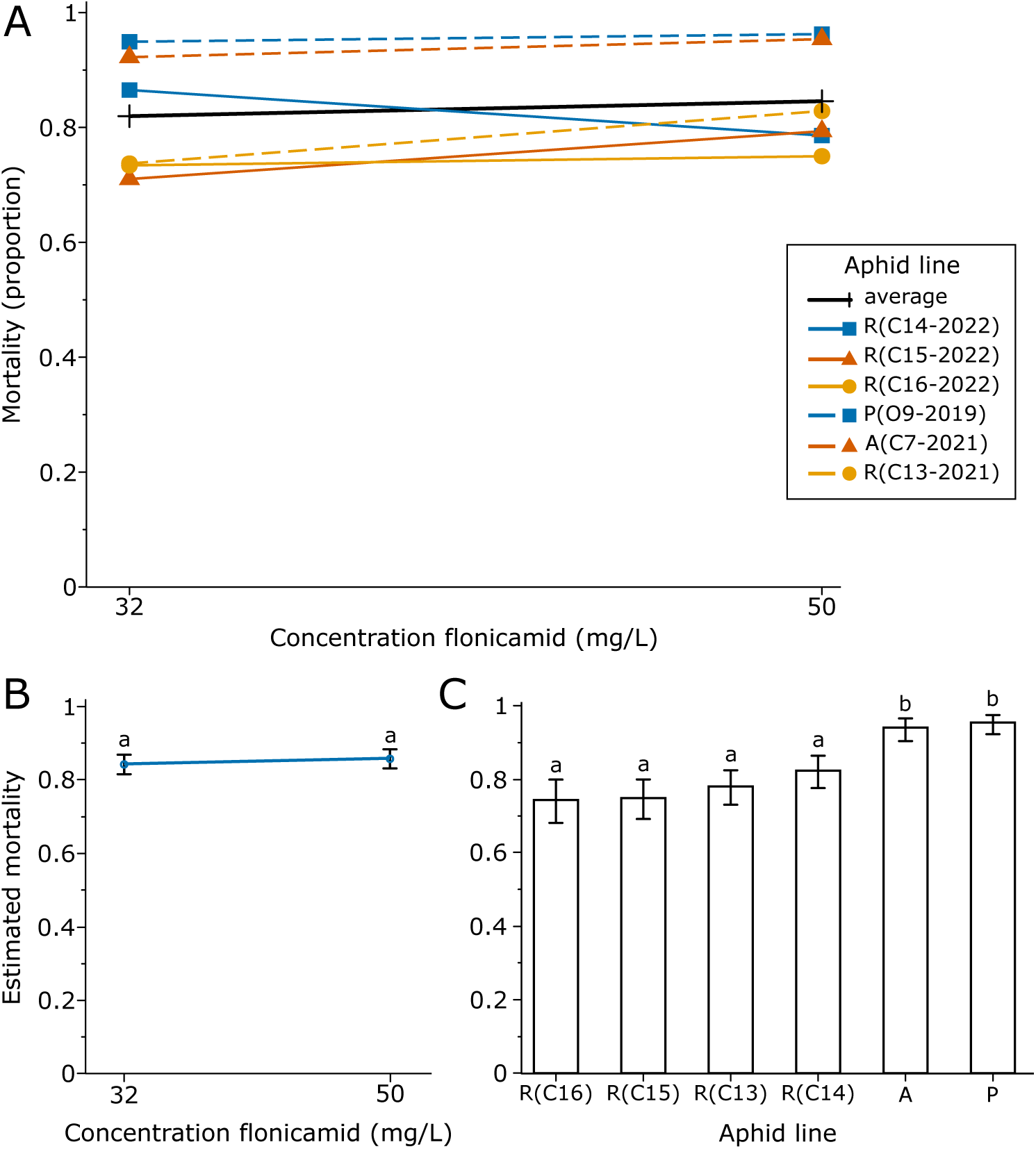
Responses of *Myzus persicae* to flonicamid. A) Average measured mortality of four aphid lines with MLG-R (R(C13-2021), R(C14-2022), R(C15-2022), and R(C16-2022)) and two lines with alternative multilocus genotypes (P(O9-2019) and A(C7-2021)) over three replicates, when exposed for 168 h to 32 and 50 mg/L flonicamid. B) Effects plot displaying the estimated effects of insecticide concentration and C) aphid line, ordered from lowest to highest average measured mortality, on aphid mortality, based on a generalised linear model. The x-axes of panels A and B are displayed on a logarithmic (base 2) scale. The error bars in panels B and C represent the standard error while the letters signify significant differences (*p <* 0.05) between treatments, as determined by Tukey’s post hoc testing. Aphid line IDs in panel C are abbreviated using the respective MLG of the line, and the greenhouse ID for lines with identical MLGs.

The observed mortality, averaged over the three replicates, ranged from 71.0% (± 3.5) for line R(C15-2022) at 32 mg/L flonicamid, to 96.3% (± 3.6) for line P(O9-2019) at 50 mg/L flonicamid (Figure 3A; Suppl. Table S6). There was no effect of insecticide concentration, 32 versus 50 mg/L flonicamid, on the mortality rate of *M. persicae* (GLM: *χ*^2^(1, N = 36) = 0.709, *p* = 0.400; Figure 3B), but there was an effect of aphid line (GLM: *χ*^2^(5, N = 36) = 88.825, *p <* 0.001; Figure 3C). Mortality of the four MLG-R lines was lower than that of the two non-MLG-R lines but did not differ within MLG (for results of Tukey’s post hoc tests, see Suppl. Table S7). The average mortality of the MLG-R lines was 77.6% (± 8.5) compared to 94.7% (± 5.2) for the non-MLG-R lines.

Additionally, we noted clear differences in body sizes between surviving aphids of MLG-R versus non-MLG-R. While the MLG-R aphids had grown during the experiment, the surviving non-MLG-R aphids were very small and did not seem to have developed past the first instar. However, these size differences were not consistently quantified as part of the experiment.

### Presence and prevalence of known insecticide resistance-inducing mutations

#### S431F (carbamates)

Mutation S431F was detected in 59% (16/27) of the MLGs (Table 4; see Suppl. File S2 for the gel electrophoresis photos). Among these are MLG-A and MLG-R, the MLGs that were dominating conventional sweet pepper greenhouses in 2019-2022 (Beekman et al., 2024). Zygosity was determined exclusively for the whole genome sequenced MLGs, revealing that all five lines carried the mutation in a heterozygous state.

**Table 4.** The genotypes of 27 different multilocus genotypes (MLGs) of *Myzus persicae* from Dutch sweet pepper, for mutations involved in resistance to carbamates (S431F) and pyrethroids (M918L/T/I, L932F, F979S, L1014F), and the genotypes of nine of these MLG for mutations associated with resistance to neonicotinoids (R81T) and tetronic and tetramic acid derivatives (A2226V). Gray fields signify mutant genotypes (M) while white fields signify the wild-type (WT). The ‘?’ represents that the zygosity of this MLG for this mutation is unknown. For M918; L = Leucine (TTG), L2 = Leucine (CTG), I = Isoleucine (ATA), and T = Threonine (ACG). The * indicates a failed PCR reaction and ** indicates failed reverse strand sequencing.

| MLG | S431F | M918 | L932F | F979S | L1014F | R81T | A2226V |
| --- | --- | --- | --- | --- | --- | --- | --- |
| A | WT/M | WT/M (L) | WT/WT | WT/WT | WT/WT | WT/WT | WT/WT |
| A2 | ?/M | WT/M (L) | WT/WT | WT/WT | WT/WT | - | - |
| B | WT/M | WT/WT | WT/WT | WT/WT | WT/WT | WT/WT | WT/WT |
| C | ?/M | WT/M (L) | WT/WT | WT/WT | WT/WT | - | - |
| D | WT/M | WT/M (T) | WT/WT | WT/WT | WT/M | WT/WT | WT/WT |
| E | WT/WT | ** | WT/WT | WT/WT | WT/WT | - | - |
| F | ?/M | WT/WT | WT/M | WT/WT | WT/M | - | - |
| G | WT/WT | WT/WT | WT/M | WT/WT | WT/M | - | - |
| H | ?/M | WT/M (L) | WT/WT | WT/WT | WT/WT | - | - |
| I | ?/M | WT/M (L<sub>2</sub>) | WT/WT | WT/WT | WT/WT | - | - |
| J | WT/WT | WT/WT | WT/M | WT/WT | WT/M | WT/WT | WT/WT |
| K | WT/WT | WT/WT | WT/M | WT/WT | WT/M | WT/WT | WT/WT |
| L | ?/M | WT/WT | WT/WT | WT/M | WT/M | - | - |
| M | ?/M | WT/WT | WT/WT | WT/WT | WT/WT | - | - |
| N | ?/M | WT/WT | WT/M | WT/WT | WT/M | - | - |
| O | WT/WT | WT/M (L) | WT/WT | WT/WT | WT/WT | - | - |
| P | WT/M | WT/M (T) | WT/WT | WT/WT | WT/M | WT/WT | WT/WT |
| P2 | ?/M | * | * | * | * | - | - |
| P3 | ?/M | WT/M (T) | WT/WT | WT/WT | WT/M | - | - |
| Q | WT/WT | WT/WT | WT/M | WT/WT | WT/M | - | - |
| R | WT/M | M/M (I/L) | WT/WT | WT/WT | WT/WT | WT/M | WT/WT |
| S | WT/WT | WT/WT | WT/WT | WT/WT | WT/WT | - | - |
| T | WT/WT | WT/WT | WT/WT | WT/WT | WT/WT | WT/WT | WT/WT |
| U | WT/WT | WT/WT | WT/WT | WT/WT | WT/WT | - | - |
| V | ?/M | ** | WT/WT | WT/WT | WT/WT | - | - |
| W | WT/WT | WT/WT | WT/WT | WT/WT | WT/WT | - | - |
| X | WT/WT | M/M (L/T) | WT/WT | WT/WT | WT/M | WT/WT | WT/WT |

#### Mutations in para (pyrethroids)

Mutations were found at all four previously known mutating sites within the investigated 578 bp stretch of *para*, the target-site for pyrethroids: L1014 (also known as *kdr*), M918 (also known as s-*kdr*), L932, and F979 (Table 4). We were unable to obtain results for MLG-P2 as PCR reactions consistently proved unsuccessful for this specific sample. At location L1014, the WT sequence is CTC. The mutant allele, TTC, substituting leucine for phenylalanine, was detected in 42% (11/26) of the MLGs. All carried a single mutant allele (henceforth called ‘heterozygous mutant’). At M918, the WT sequence is ATG which codes for methionine. The detected mutant sequences either cause a substitution of methionine into leucine (TTG; 7/24, CTG; 1/24), isoleucine (ATA; 1/24), or threonine (ACG; 4/24). For this location, 46% (11/24) of the MLGs carry at least one mutant allele. For MLG-E and MLG-V, sequencing of the reverse strand consistently failed, preventing us from determining the genotype for these MLGs at location 918. At L932, the WT sequence is TTG. Mutants are heterozygous for a substitution of leucine into phenylalanine (TTT). This mutation was found in 23% (6/26) of the MLGs. Mutation F979S (TTT; phenylalanine to TCT; serine) was only found once, in MLG-L, which is heterozygous for this mutation.

The wild-type (WT) genotype for the investigated region of *para*, at all four mutating locations, was detected in 25% (6/24) of the MLGs. Additionally, also MLG-E and MLG-V have the WT alleles at the three determined locations, although their genotype at location 918 is unknown. In 29% (7/24) of the MLGs, mutations are only present at M918, and all these MLGs substitute methionine for leucine. All MLGs with M918T (17%; 4/24), L932F (23%; 6/26) or F979S (4%; 1/26) also carry the L1014F substitution. All MLGs with a mutation at location 918 are heterozygous mutants, except for MLG-R (TTG/ATA) and MLG-X (ACG/TTG) which both contain two different mutant alleles (henceforth called ‘transheterozygous mutant’). No MLGs were found with mutations at more than two locations of the gene.

#### R81T (neonicotinoids) and A2226V (tetronic and tetramic acid derivatives)

We did not manage to reliably detect mutation R81T using PCR. Therefore, we were only able to search for this mutation in the genomes of the whole genome sequenced MLGs (MLG-A, B, D, J, K, P, R, T, and X). Simularly, mutation A2226V was only searched for in the MLGs with available genomes. Mutation R81T was found heterozygously only in MLG-R, while mutation A2226V was not detected in any of these MLGs (Table 4).

## Discussion

The green peach aphid *M. persicae* is notorious for its ability to quickly become resistant to insecticides. Regular monitoring of populations of *M. persicae* for resistance becomes increasingly important as the selection of effective insecticides, especially those compatible with biocontrol agents, diminishes. Up to now, no resistance has been reported for *M. persicae* to the insecticides pymetrozine and flonicamid, which are compatible with non-hemipteran biocontrol agents (Mota-Sanchez and Wise, 2024). However, our previous work on the genetic structure of this population showed strong dominance of specific MLGs under the application of these insecticides (Beekman et al., 2024). This suggests positive selection, potentially due to reduced sensitivity to the respective insecticides. Therefore, we tested various lines of *M. persicae* from sweet pepper for their sensitivity to pymetrozine and flonicamid, among which the lines with the dominant MLGs. In brief, we found that the dominant MLGs under the application of either flonicamid or pymetrozine were the least sensitive to the respective insecticides, indicating that sensitivity to these insecticides could underlie clonal selection at the greenhouse scale.

Additionally, we examined the prevalence of various mutations associated with insecticide resistance in 27 MLGs of the *M. persicae* population collected from Dutch sweet pepper greenhouses between 2019 and 2022. Our findings revealed a high prevalence of mutation S431F, which confers resistance to carbamates, as well as a high prevalence of mutations in *para*, conferring resistance to pyrethroids. Furthermore, mutation R81T, linked with resistance to neonicotinoids, was detected once, while mutation A2226V, involved in resistance to tetronic and tetramic acid derivatives, was not detected.

### Sensitivity to pymetrozine and flonicamid

The results of the pymetrozine and flonicamid assays indicate that the MLGs of *M. persicae* that dominated conventional Dutch sweet pepper greenhouses in 2019 and 2022 exhibit decreased susceptibility to these insecticides, when compared to the sensitivity of other MLGs. Importantly, quantification of the actual levels of resistance was not the goal of this study, and was also not possible due to two main factors: firstly, we did not obtain 100% mortality across all aphid lines, which hindered dose-response analyses and the reliable calculation LC_50_-values; secondly, the absence of a susceptible *M. persicae* line, necessary for determining resistance ratios (RRs), which are calculated by dividing the LC_50_-value of a studied line by the LC_50_-value of a susceptible line. Resistance ratio’s are translated to resistance levels in the following way: susceptible (RR ¡ 5-fold), low resistance (RR = 5 to 10-fold), moderate resistance (RR = 10 to 40-fold), high resistance (RR = 40 to 160-fold), and extremely high resistance (RR ¿ 160-fold) (Organization, 1980). While previous studies have investigated RRs of field populations of *M. persicae* to pymetrozine and flonicamid, none have linked the sensitivity of the respective aphid lines to their prevalence in the field, as demonstrated in this study (Foster et al., 2002; Hu et al., 2023; Margaritopoulos et al., 2010; Margaritopoulos et al., 2021).

#### Pymetrozine

Earlier studies reported maximum RRs for *M. persicae* populations from Greece and England of around RR = 6, indicating no or very low levels of resistance and little variation in sensitivity between the tested lines (Foster et al., 2002; Margaritopoulos et al., 2010). In comparison, our results display a wide range of susceptibility levels. Our most sensitive lines displayed higher levels of mortality at 2 mg/L than our least sensitive lines did at 40 mg/L. The mortality rates we observed for the most sensitive lines closely resemble the LC_90_ of 6.92 mg/L after 96 h exposure reported for *M. persicae* from the North Western Himalayan region (Paschapur et al., 2019). This suggests that our least sensitive lines, MLG-A and MLG-R, are less susceptible to pymetrozine than the North Western Himalayan population. Recently, a much higher RR (34.8) was reported for a population of *M. persicae* from cabbage collected from the southeastern coast of China (Hu et al., 2023). They observed an LC_50_-value of 0.86 mg/L for the susceptible line and 29.86 mg/L for the field population after four days of exposure. These results show similarities to ours, with our most sensitive line displaying on average over 50% mortality at 2 mg/L, and our least sensitive line reaching 50% mortality on average somewhere between 10 and 40 mg/L.

#### Flonicamid

The first reported LC_50_ for *M. persicae* exposed to flonicamid was 0.61-0.77 mg/L after five days of exposure (Morita et al., 2007). Recent studies have reported RRs for *M. periscae* of around five to six, for populations from China and Greece (Hu et al., 2023; Margaritopoulos et al., 2021). Another recent study, exposing *M. persicae* to flonicamid for as long as 168 h, similar as we did in experiment F2, found no differences in sensitivity between various populations from Australian agricultural populations, primarily from *Brassica napus* (Arthur et al., 2022). While our study did not yield RRs, the recommended field dose of 50 mg/L flonicamid only resulted in approximately 78% mortality of MLG-R aphids, implying that aphids with this genotype can survive conventional flonicamid treatments in Dutch sweet pepper greenhouses. The effects of flonicamid treatments in a greenhouse likely differ from those observed in the bioassays, given that aphids in greenhouses are exposed to multiple stressors simultaneously, unlike the single stressor administered in our study. However, our results probably explain the extreme difficulties faced in controlling *M. persicae* in conventional Dutch sweet pepper greenhouses, as reported in autumn 2021 and spring 2022 (Biobest Group, 2022; Glastuinbouw Nederland, 2022). The high selection pressure exerted by flonicamid likely has selected for *M. persicae* genotypes capable of surviving and reproducing at the recommended field dose, rapidly leading to the development of decreased sensitivity or even resistance.

#### Extended exposure for full effects

Given the mode of action of pymetrozine and flonicamid as selective feeding blockers, and their slow-acting nature, IRAC recommends assessing mortality after 120 h of exposure, although it is not specified whether this refers to 120 h after the placement of adults or after the nymphs are born. Due to logistical constraints, we concluded our final mortality assessment one day earlier than IRAC recommendations, at 96 h after introduction of the adults. At this point, it is possible that not all affected aphids had died yet. Arthur et al. (2022) even demonstrated that exposure of *M. persicae* to flonicamid for 144 h or longer may be necessary to capture the full effects. Furthermore, our highest tested concentrations in experiments PF1, based on concentrations used in previous studies (Cho et al., 2011; Foster et al., 2002; Margaritopoulos et al., 2010; Morita et al., 2007), were lower than those recommended for aphid control in Dutch sweet pepper greenhouses. The recommended field concentration for pymetrozine was 100 mg/L (Plenum 50 WG, instruction manual W2; Ctgb, 2024), and is 50 mg/L for flonicamid (Teppeki WG, instruction manual W4; Ctgb, 2024). Only during experiment F2 did we test the recommended field concentration, and did we expose the aphids for 168 h. Nevertheless, consistent 100% mortality was not achieved. However, mortality of line R(C13-2021) increased significantly with the extended exposure time, from 37 % to 74%, averaged over the three replicates, after 96 h and 168 h exposure respectively, making it evident that 96 h of exposure is insufficient to fully capture the effects of flonicamid.

#### Sublethal effects

During experiment F2, we observed notable differences in body size among surviving aphids of different MLGs. Some aphids that survived the insecticide treatment up until 168 h post-initial adult placement seemed to have not developed beyond the first instar. Consequently, the differences in effect sizes of insecticide treatment between different MLGs are expected to be more pronounced on a community scale, as aphids that fail to develop will not contribute to population growth. Delayed development as a consequence of flonicamid exposure has been demonstrated before, already at concentrations of 1.25 mg/L and below, both for *M. persicae* (Cho et al., 2011) and *A. gossypii* (Koo et al., 2015). To address these factors in future experiments, we recommend conducting mortality assessments even later, measuring total population growth, or including observations of aphid developmental stages.

#### Potential resistance through metabolic detoxification

Resistance to pymetrozine has been observed in various non-aphid insect species (W. Li et al., 2020; Song et al., 2022; Wang et al., 2021; Y. Zhang et al., 2017). This resistance has been linked to the upregulation of various genes encoding cytochrome P450 enzymes responsible for pymetrozine metabolism (Gong et al., 2023; Nauen et al., 2013; Wang et al., 2021; Y. Zhang et al., 2017). Similarly, resistance to flonicamid has been documented in the cotton aphid *A. gossypii* (Koo et al., 2014; Shi et al., 2023), and for various other non-aphid insect species (Abbas et al., 2021; Roy et al., 2019; Ullah et al., 2021; T.-y. Zhang et al., 2023). Although the mechanisms of flonicamid resistance remain unclear, overexpression of cytochrome P450 genes has also here been implicated in reducing sensitivity in the red imported fire ant and in *D. melanogaster* (T. Zhang et al., 2023). However, honey bees have demonstrated a different resistance mechanism to flonicamid. They typically encode *Naam2*, an enzyme that is involved in a pathway leading to the biosynthesis of NAD+ (nicotinamide adenine dinucleotide), and is inhibited by a metabolite of flonicamid (Qiao et al., 2022). A recombinant form of *Naam2* circumvents this inhibition and confers resistance to flonicamid.

The upregulation of a cytochrome P450 gene, specifically *CYP6CY3*, has also been documented in *M. persicae* and is known to contribute to resistance to neonicotinoids (Bass et al., 2011; Philippou et al., 2010; Puinean et al., 2010; Singh et al., 2021) and sulfoxaflor (Pym et al., 2022). To investigate whether cytochrome P450 enzymes play a role in the decreased sensitivity of MLG-A and MLG-R to pymetrozine, and MLG-R to flonicamid, future research should explore the effects of adding piperonyl butoxide (PBO), a P450 enzyme inhibitor, to the insecticide sensitivity assays. Alternatively, genome-wide association studies, including the genomes of MLG-A and MLG-R, may elucidate potential mutations leading to resistance to the insecticides.

### Detection of known mutations

#### S431F (carbamates)

The detected frequency of mutation S431F in 59% (16/27) of the MLGs is close to the 53% (67/127) that was determined for global *M. persicae* populations by Singh et al. (2021). Specifically, they found S431F in 80% (8/10) of the aphids lines collected from pepper plants, whereas a study in sweet pepper fields in Central Chile reported only an average 36% prevalence of the mutation over four populations which comprised 202 MLGs in total (Rubiano-Rodríguez et al., 2014). While we could not assess the zygosity of this mutation for all MLGs, all five whole-genome-sequenced MLGs carrying the S431F mutation were heterozygous, confirming previous studies (Anstead et al., 2008; Fontaine et al., 2011; Mingeot et al., 2021). Although S431F homozygotes are occasionally observed, primarily in holocyclic populations (Charaabi et al., 2016; Criniti et al., 2008; Rubiano-Rodríguez et al., 2014; Voudouris et al., 2016), the high prevalence of S431F heterozygotes may be linked to observed fitness costs associated with this mutation in *M. persicae* (Foster et al., 2003), which suggests especially high fitness costs of the homozygous form. Heterozygous S431F mutants of *M. persicae* can display ¿100 fold insensitivity to the dimethyl carbamate pirimicarb (Moores et al., 1994). Pirimicarb, despite being less effective against *M. persicae*, is commonly utilised in Dutch sweet pepper greenhouses primarily for the control of other aphid species such as *Aulacorthum solani* (Beekman et al., 2022; Van Iperen, 2024). Our data indicates a high prevalence of S431F in *M. persicae* from Dutch sweet pepper, notably also in MLG-A and MLG-R that were dominant in Dutch sweet pepper greenhouses in 2019 and 2022 respectively (Beekman et al., 2024). This high prevalence of S431F explains why pirimicarb often fails to control *M. persicae* populations in this crop system.

#### Mutations in para (pyrethroids)

Mutations in *para*, the target-site for pyrethroids, were prevalent in our study. The WT genotype for the studied stretch of *para* was detected in 25% (6/24) of the MLGs. MLG-E and MLG-V exhibited WT alleles at the three determined locations, while their genotype at location 918 remains unknown. All other MLGs carried mutations at one or multiple locations: M918, L932, F979, and L1014.

We found mutation L1014F, known as *kdr*, in 41% (11/27) of the MLGs, all of which were heterozygous mutants. In previous studies, L1014F was detected in only 20% (2/10) of global *M. persicae* lines from pepper (Singh et al., 2021), and in 69% of MLGs from a sweet pepper field for Chile, with 13% being homozygous mutants (Rubiano-Rodríguez et al., 2014). The frequency we observed thus lies in between the large range of the previous findings. Although no homozygous L1014F mutants were found in our study, they have been reported occasionally in other studies, albeit less frequent than heterozygous forms (Charaabi et al., 2016; Eleftherianos et al., 2008; Hlaoui et al., 2022).

At location M918, known as s-*kdr*, four different substitutions were detected: methionine (ATG) into leucine (TTG, CTG), isoleucine (ATA), or threonine (ACG). M918 mutations were consistently found in the heterozygous state, except for two transheterozygous mutants: MLG-X (ACG/TTG) and MLG-R (TTG/ATA). The combination ACG/TTG was previously identified by Panini et al. (2015), demonstrating that M918 transheterozygous aphids, carrying both M918L and M918T, had increased resistance levels to both type I and type II pyrethroids compared to either mutation alone. M918L combined with M918I (TTG/ATA) was first described for *M. persicae* in a sample from China by Singh et al. (2021), who argued that isoleucine at location M918 makes the sodium channel more ‘mammalian-like’, reducing sensitivity to insect-specific pyrethroids. We found M918T consistently together with L1014F, in line with previous literature (Bass et al., 2014; Panini et al., 2015). This combination of mutations is reported to result in approximately a 10x increase in pyrethroid resistance compared to L1014F alone (as summarised by Soderlund, 2008). In addition, we frequently detected M918L without L1014F, consistent with prior studies (Fontaine et al., 2011; Panini et al., 2014; Singh et al., 2021). M918L has been shown to confer strong resistance to both type I and II pyrethroids in *M. persicae* (Fontaine et al., 2011; Hlaoui et al., 2022).

For mutation L932F, 22% (6/27) of the MLGs were heterozygous. Additionally, we found that all MLGs carrying this mutation also harboured L1014F, a linkage also observed in 100% (16/16) *M. persicae* from oilseed rape in France that carried L932F (Fontaine et al., 2011) and in 91% (10/11) of *M. persicae* samples carrying L932F from oilseed rape in Czechia (Stará et al., 2024). However, mutation L932F is seldom reported, and its effects on *M. persicae* susceptibility to pyrethroids are still unclear.

Mutation F979S is also rarely reported in the literature. We detected it once in MLG-L, which is heterozygous for this mutation and also carries L1014F. Previous studies have also found F979S linked with L1014F (Cassanelli et al., 2005; Criniti et al., 2008). Mutation F979S in *M. persicae* potentially has an additive effect on levels of pyrethroid resistance caused by L1014F (Criniti et al., 2008).

#### R81T (neonicotinoids)

We detected mutation R81T in one of the nine genomes, in MLG-R, in the heterozygous state. To our knowledge, this observation represents the northernmost occurrence of this mutation, with the previous northernmost detection being in Belgium (IRAC, 2019). R81T was initially found in a *M. persicae* clone from France in 2009 (Bass et al., 2011) and is mainly distributed in Europe around the Mediterranean, including France, Italy, Spain, Greece (Mezei et al., 2022; Singh et al., 2021; Slater et al., 2012; Voudouris et al., 2017), Tunisia (Charaabi et al., 2018), and Morocco (IRAC, 2019), but has also been detected in China (Xu et al., 2022). Interestingly, Singh et al. (2021) found R81T exclusively in clones overexpressing the specific cytochrome P450 gene *CYP6CY3*, which encodes an enzyme known to be involved in metabolic detoxification of neonicotinoids (Bass et al., 2013; Bass et al., 2011; Philippou et al., 2010; Puinean et al., 2010). Although not examined in this study, it is plausible that MLG-R also overexpresses *CYP6CY3*.

It is possible that some of our MLGs with no genome sequences available also carry the R81T mutation. However, a previous study, analysing the genomes of *M. persicae* from around the world, detected R81T in only 17 out of the 127 analysed clones (Singh et al., 2021), emphasising its rarity. We attempted to detect R81T in all MLGs but were unable to obtain clear PCR results using the methods described by Panini et al. (2014). Upon detailed examination of the *M. persicae* genome, we discovered that the target site of primer MpNACR-FW did not match the reference sequence. According to our sequenced *M. persicae* lines and to the variant call format (vcf) file kindly provided by Singh et al. (2021), which includes genomes from 127 worldwide collected *M. persicae* lines, we suggest the correct sequence for this primer to be 5’-**<u>A</u>**ATAATGAAATCAAACGTTTGGTTGAG-3’. Similarly, the sequence of primer MpNACR-R514 was also found incorrect and should be 5’-GAGA**<u>A</u>**AAATCGCTGAGTAGATTTC-3’. Future research can test the efficacy of these arguably improved primers.

#### A2226V (tetronic and tetramic acid derivatives)

The A2226V mutation, linked to resistance to tetronic and tetramic acid derivatives, was not found in any of the whole genome sequenced *M. persicae* lines. Although it is possible that this mutation is present in some of our unanalysed MLGs, its has currently only been detected in samples from Australia (Singh et al., 2021; Umina et al., 2022).

#### One sample tested per MLG

We examined the resistance genotype of the MLGs for S431F and *para* in only one sample per MLG, except for MLG-A, where 13 samples were checked. Our results showed consistent resistance genotypes in all 13 samples of MLG-A. However, previous studies have occasionally observed varying resistance genotypes among *M. persicae* samples with the same MLG (Mingeot et al., 2021; Zamoum et al., 2005). This variability could especially be expected in MLGs with high prevalence and widespread distribution, where differences in environmental pressures between habitats can select for variable resistance genotypes. Several *M. persicae* MLGs investigated in this study were originally detected in various greenhouses (Beekman et al., 2024), sometimes in both greenhouses with conventional pest management and those with organic pest management, where environmental selection for mutations involved in insecticide resistance could have differed. Therefore, it is possible that some unique resistance genotypes were missed in our study by analysing only one sample per MLG.

### Future control of *Myzus persicae*

Combining the detected mutations with the results of the insecticide sensitivity assays (see summarising Figure 4) reveals that MLG-A remains susceptible to flonicamid, and likely also to spirotetramat (belonging to the tetronic and tetramic acid derivatives, applied in greenhouses against aphids) and neonicotinoids, although we did not test for resistance through metabolic detoxification, which is also involved in resistance to the latter group of insecticides. Conversely, for MLG-R, our findings suggest a more concerning outlook as this MLG exhibited reduced sensitivity to both pymetrozine and flonicamid, and harbours the mutations that are linked to resistance to carbamates, pyrethroids, and neonicotinoids. Of the insecticides we investigated, only spirotetramat, which is currently banned in the EU (Ctgb, 2024), would be able to control MLG-R. The accumulation of resistance to various insecticidal groups in a single aphid genotype likely causes these genotypes to be highly successful in conventional agricultural systems, posing continuous challenges to growers in the control of these genotypes with insecticides.

**Figure 4.**
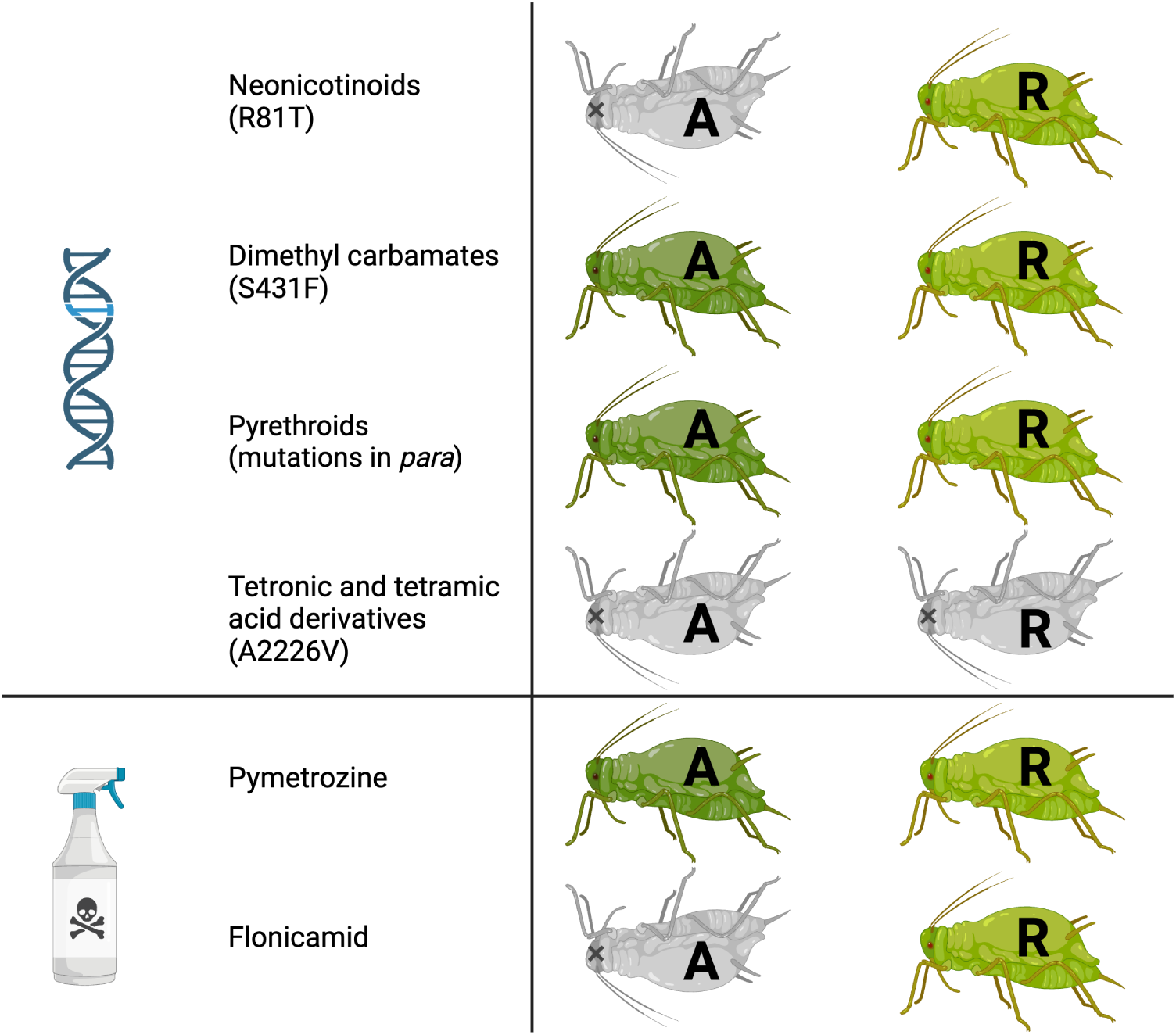
Summary of the observed susceptibility of two multilocus genotypes (MLGs), MLG-A and MLG-R, of *Myzus persicae* that dominated in Dutch sweet pepper greenhouses during 2019/2020 and 2021/2022 respectively (Beekman et al., 2024), to various insecticides. We determined the susceptibility to neonicotinoids, dimethyl carbamates, pyrethroids, and tetronic and tetramic acid derivatives by genetic screening for known insecticide resistance mutations. We assessed susceptibility to pymetrozine and flonicamid through insecticide sensitivity assays. Grey upside-down aphids indicate susceptibility of the genotype to the insecticide, while green upright aphids indicate resistance or reduced susceptible to the insecticide. The letter (A/R) inside the aphid indicates the MLG of the aphid line. Figure created with BioRender.com.

Since mid-2022, cyantraniliprole, a new insecticide targeting insect ryanodine receptors (Jeanguenat, 2013), has been approved for controlling *M. persicae* in Dutch sweet pepper (Ctgb, 2024). This insecticide has effectively managed *M. persicae* populations in conventional Dutch sweet pepper greenhouses in 2022 and 2023, which were previously challenging to control chemically (Jeannette Vriend, Glastuinbouw NL, personal communication). As of now, no resistance to cyantraniliprole has been reported in *M. persicae* (Mota-Sanchez and Wise, 2024). However, various missense mutations have already been linked to target site insensitivity in various Lepidopteran species, leading to moderate to high levels of resistance (Jouraku et al., 2020; Zuo et al., 2020), and upregulation of cytochrome P450 monooxygenase and UDP-glycosyltransferase genes has been shown to result in cyantraniliprole resistance in the cotton aphid *A. gossypii* (Zeng et al., 2021). Therefore, we can anticipate a significant potential for the development of resistance of *M. persicae* to cyantraniliprole.

The devastating effects of insecticides on the environment and on organismal health have become increasingly clear (Ansari et al., 2014; Geiger et al., 2010; Kim et al., 2017; Pimentel, 2005). Additionally, the number of available effective insecticidal groups is diminishing rapidly, and the discovery of new chemical groups with insecticidal activity is becoming increasingly rare. Our study, showing the prevalence of resistance to various insecticidal groups in *M. persicae* from sweet pepper greenhouses, underscores the necessity for more sustainable control strategies. Biocontrol offers an effective and sustainable alternatives to conventional insecticides in controlling *M. persicae* as a crop pest, especially in greenhouse environments (Dedryver et al., 2010; Messelink et al., 2012; Messelink et al., 2024). However, supplementing biocontrol with chemical insecticides, a common practice in many crop systems including Dutch sweet pepper, can detrimentally effects biocontrol communities (Cloyd and Bethke, 2011; Desneux et al., 2007), consequently diminishing biocontrol effectiveness or even leading to no reduction in pest population densities at all (Janssen and van Rijn, 2021). Therefore, pest management strategies that prioritise biocontrol and minimise chemical insecticide use are critical for sustainable agriculture and environmental and organismal health.

## Acknowledgements

We are very grateful to Jordy Litjens for taking care of the aphid rearing and for his help during the insecticide sensitivity assays. We would like to thank Helena Donner, Tom Groot, Gerben Messelink, Ben Philip, and Christoph Vorburger for their ideas and feedback during discussions.

## Data availability statement

The Supplementary Tables are available on figshare and are accessible via the following link: https://doi.org/10.6084/m9.figshare.26147002.

## CRediT authorship contribution statements

**Mariska M. Beekman:** Conceptualisation, Formal analysis, Funding acquisition, Investigation, Methodology, Supervision, Visualisation, Writing – original draft, Writing – review & editing. **Xinyan Ruan**: Investigation, Methodology. **Marcel Dicke**: Supervision, Writing – review & editing. **Bas J. Zwaan:** Funding acquisition, Supervision, Writing – review & editing. **Bart A. Pannebakker:** Funding acquisition, Project administration, Methodology, Supervision, Writing – review & editing **Eveline C. Verhulst:** Funding acquisition, Methodology, Supervision, Writing – review & editing.

