## Supplementary Materials for "Insecticide resistance in *Myzus persicae* collected from sweet pepper"

#### Supplementary Tables

All supplementary tables are accessible via the following link:
https://doi.org/10.6084/m9.figshare.26147002

Table S1: Locations of known mutations linked to insecticide resistance in reference genome G006 v.3 of *Myzus persicae*

Table S2: Raw data of pymetrozine experiment 1

Table S3: Results of Tukey post hoc test on the generalised linear model, effects ”aphid line” and ”insecticide concentration” - pymetrozine.

Table S4: Raw data of flonicamid experiment 1

Table S5: Results of Tukey post hoc test on the generalised linear model, effects ”aphid line” and ”insecticide concentration” - flonicamid exp. 1.

Table S6: Raw data of flonicamid experiment 2

Table S7: Results of Tukey post hoc test on the generalised linear model, effects ”aphid line” and ”insecticide concentration” - flonicamid exp. 2.

### Supplementary File S1: Check genotypes of aphids tested in the pymetrozine and flonicamid sensitivity assays

#### Methods

The genotypes were determined for three (pymetrozine and flonicamid experiment 1 (PF1)) or four (Flonicamid experiment 2 (F2)) randomly sampled aphids from each of the three control treatment at the end of each experiment. Because the MLGs for all aphid lines were known beforehand, only one or a few microsatellite markers sufficed to check the genotypes. The control samples of the pymetrozine experiment were checked with marker myz2 (Figure S1-1). Those of flonicamid experiment 1 were checked with M86 and myz9 (Figure S1-2). For experiment F2, one sample of each line was first checked with M40 and M86, and myz9, which showed that myz9 alone was enough to visualise all different MLGs (Figure S1-3).

#### Results

Experiment PF1: Aphid line T(C12-2021)*, which was expected to have MLG-A based on earlier genotyping, turned out to display the same pattern on the agarose gel as line T(C12-2021), which had MLG-T. We suspect that this aphid line has accidentally gotten mixed up with line T(C12-2021) at some time between the initial genotyping and submitting this line to the insecticide sensitivity assays. Hence, this line is treated as MLG-T in the entire manuscript.

Experiment F2: Aphid line 'bouvardia' displayed a mix of two genotypes and was thus omitted from further analyses. Line 'koppert' displayed the same genotype as P(09-2019) and thus we suspect this line had gotten mixed up due to human errors. Therefore, this line was also omitted from further analyses.

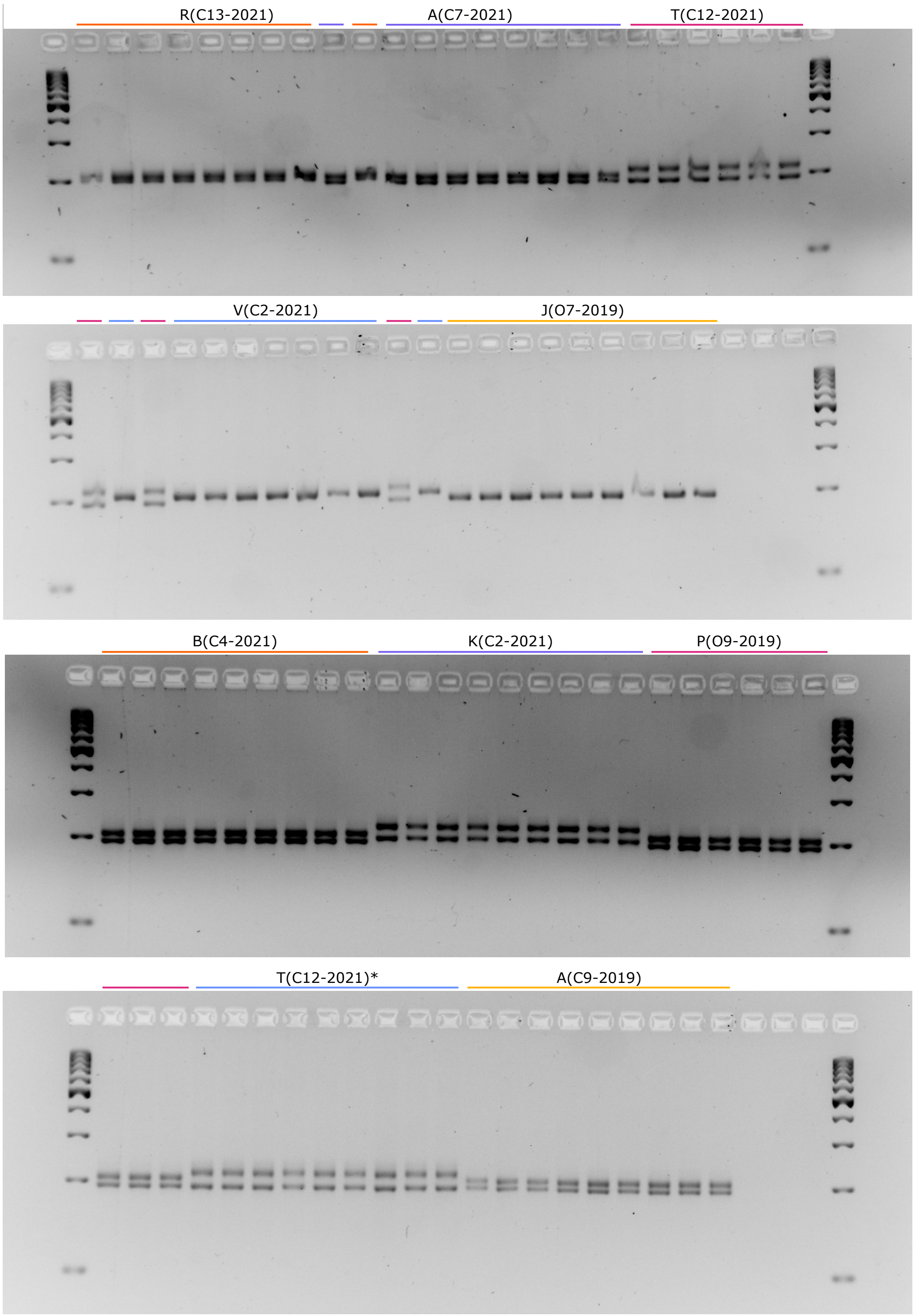


**Figure S1-1.** Genotypes check of aphids included in the pymetrozine sensitivity experiment. On each gel, the first and last wells display the GeneRuler 100bp DNA Ladder (Thermo Scientific).

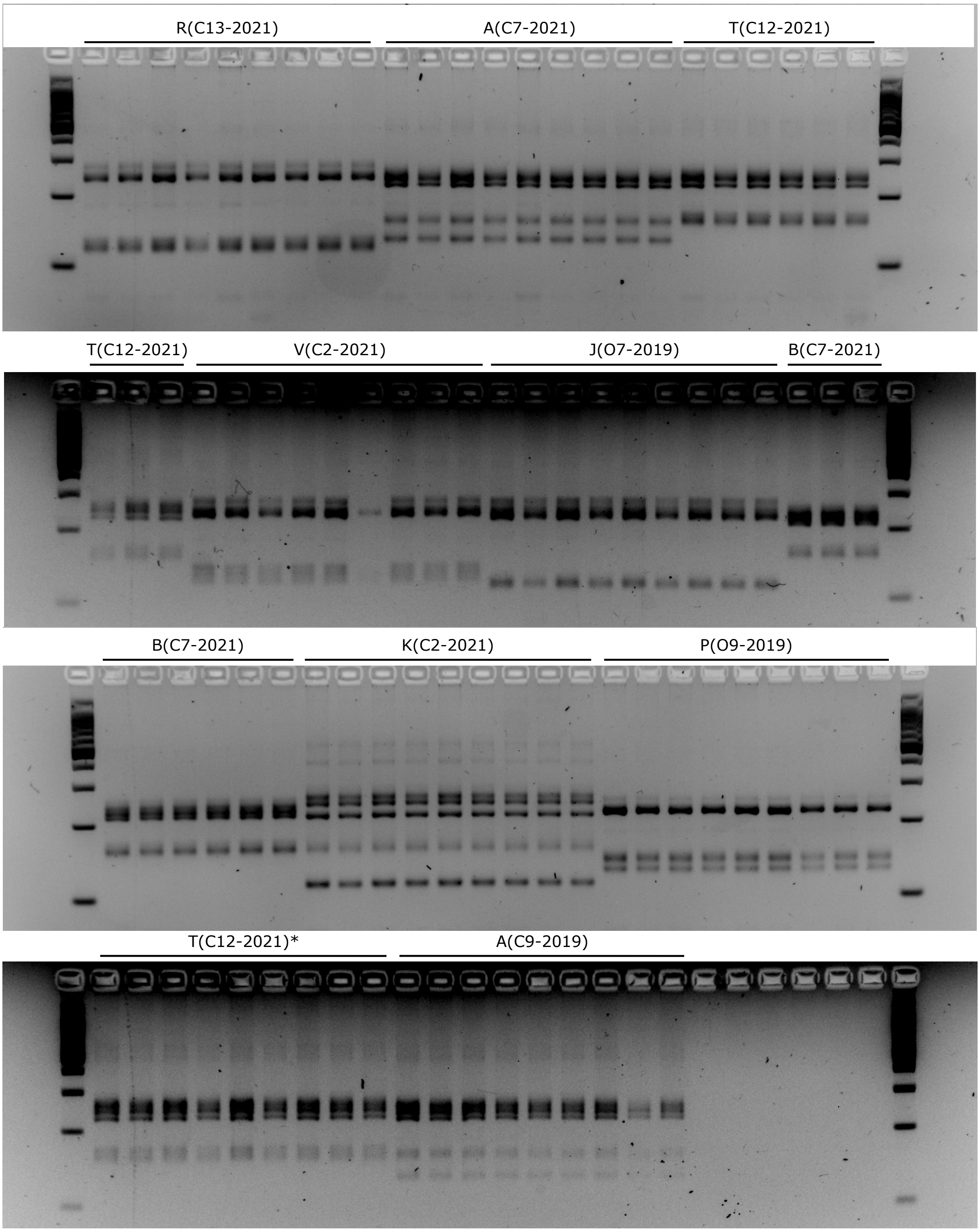


**Figure S1-2.** Genotypes check of aphids included in the flonicamid sensitivity experiment. On each gel, the first and last wells display the GeneRuler 100bp DNA Ladder (Thermo Scientific).

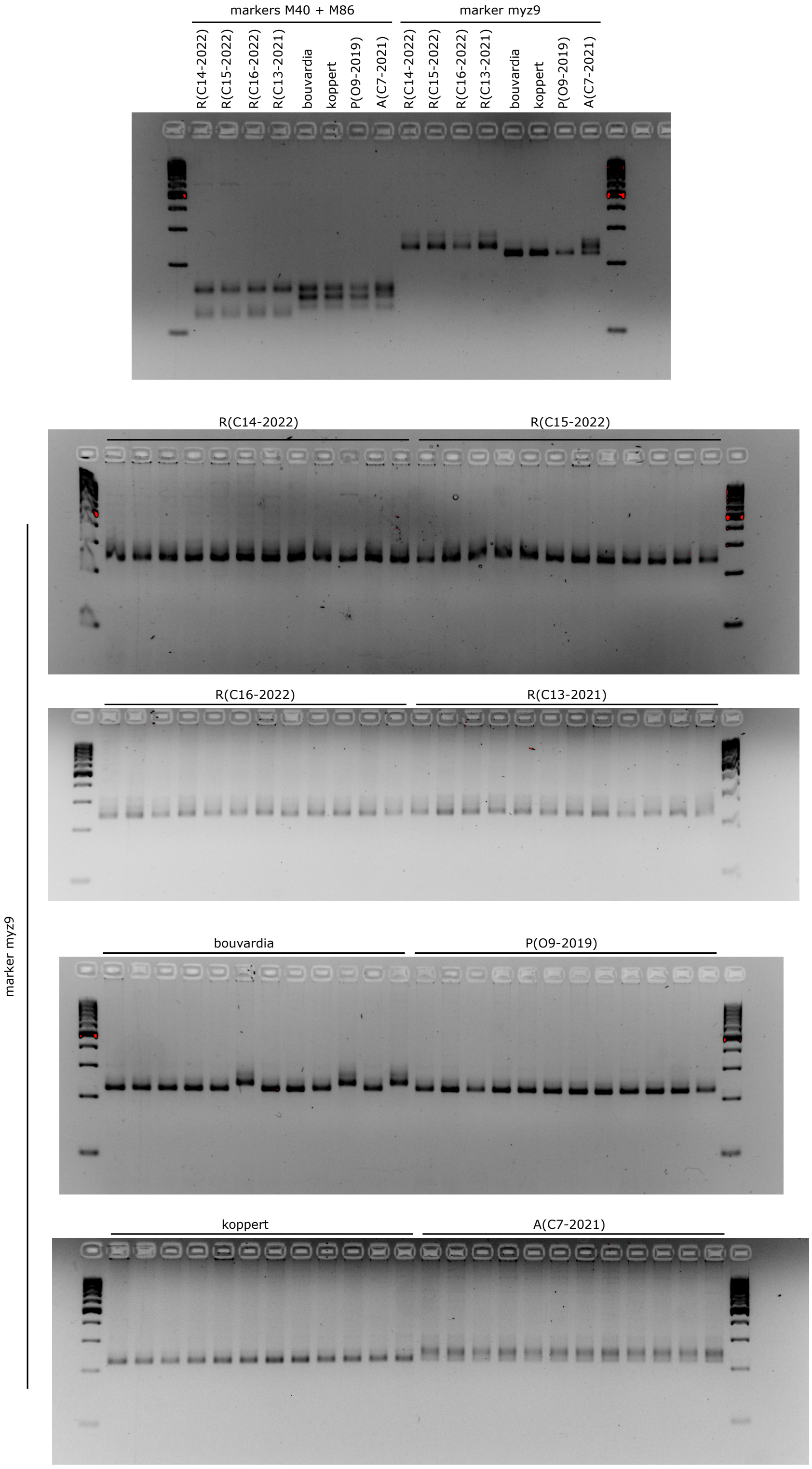


**Figure S1-3.** Genotypes check of aphids included in the flonicamid sensitivity experiment 2. On each gel, the first and last wells display the GeneRuler 100bp DNA Ladder (Thermo Scientific).

### Supplementary File S2: Gel photos detecting MACE mutation

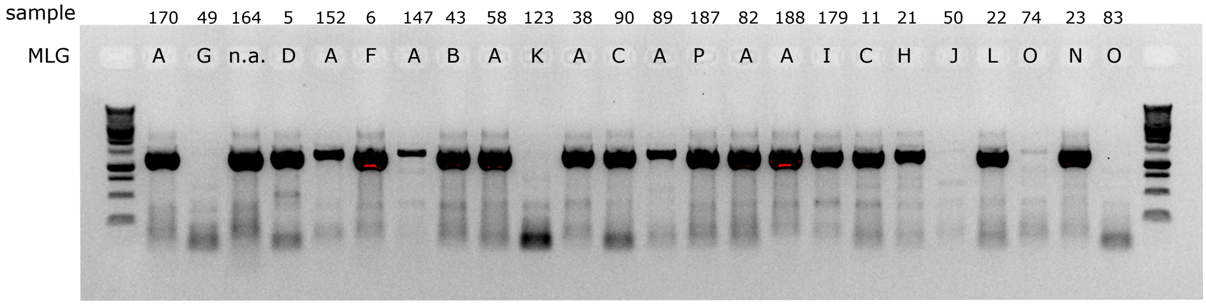


**Figure S2-1.** Detection of MACE/S431F mutation in *Myzus persicae* genotypes A-D, F-L, and N-P. The MLG of sample 164 could not be determined. The first and last well display the Promega 1kb DNA ladder.

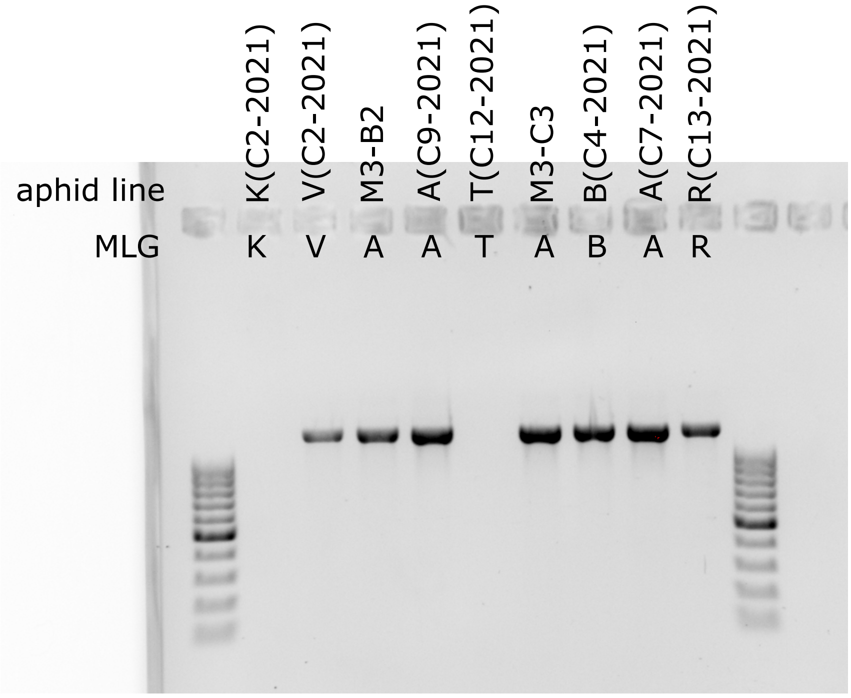


**Figure S2-2.** Detection of MACE/S431F mutation in *Myzus persicae* genotypes A, B, K, R, T, and V. The first and last well display the GeneRuler 100bp DNA Ladder (Thermo Scientific).

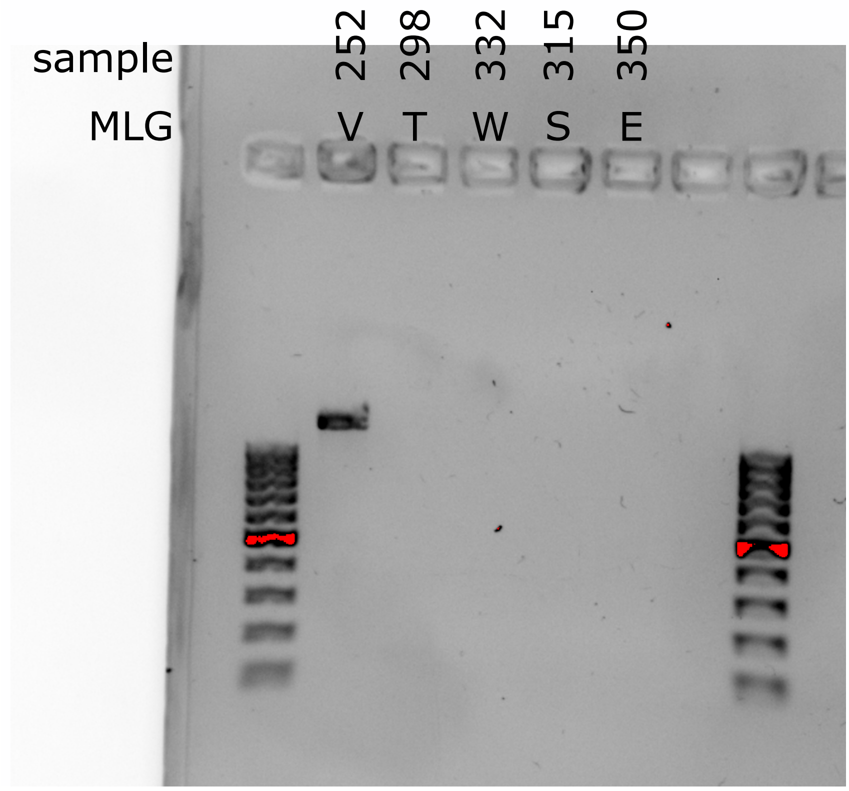


**Figure S2-3.** Detection of MACE/S431F mutation in *Myzus persicae* genotypes E, S, T, V, and W. The first and last well display the GeneRuler 100bp DNA Ladder (Thermo Scientific).

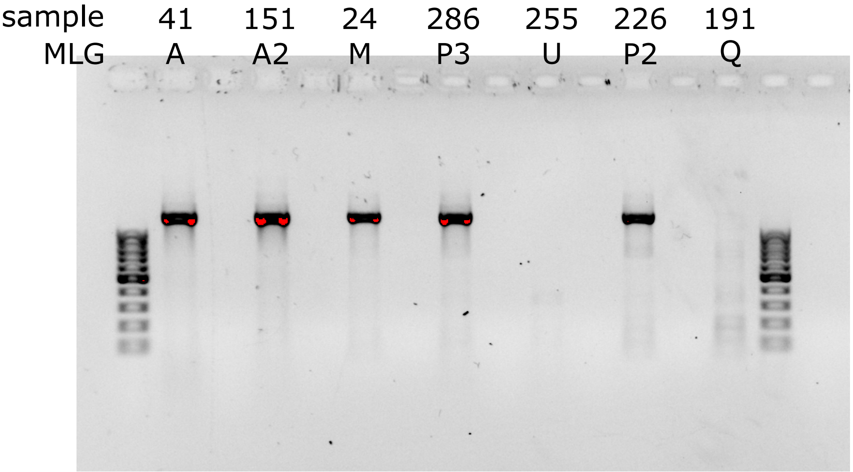


**Figure S2-4.** Detection of MACE/S431F mutation in *Myzus persicae* genotypes A, A2, M, P2, P3, Q, and U. The first and last well display the GeneRuler 100bp DNA Ladder (Thermo Scientific).

#### Supplementary File S3: Attempted optimisation of R81T detection by PCR

We attempted to detect mutation R81T with PCR as described by Panini et al. (2014). Specifically, primers MpNACR-FW and MpNACR-R514 (Table 3) are expected to amplify a 177 bp fragment of the WT allele, while MpNACRr-RE and MpNACR-F52 should amplify a 332 bp fragment of the mutant allele. Together, primers MpNACR-R514 and MpNACR-F52 are anticipated to amplify a 508 bp fragment, common to both the WT and mutant alleles, serving as a control.

##### Trial 1

Using GoTaq polymerase and buffer, equal volumes of primers.

PCR program: 94℃ for 2min, 30 cycles of 94℃ for 30s, 48℃ for 30s and 72℃ for 45s. Final extension 72℃ for 5min.

Results: no control fragment. There is a bright 200 bp fragment and sometimes a very faint band which could be 177 bp.

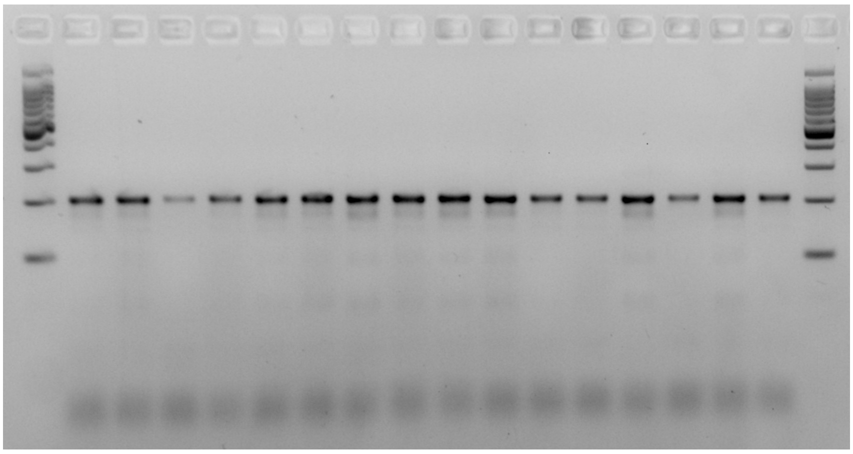

##### Trial 2

Switching to Qiagen Multiplex PCR mix, testing three annealing temperatures, equal volume of primers.

PCR program: 95℃ 15min, 35 cycles of 94℃ for 30s, 50/ 55/ 60℃ for 45s, 72℃ for 45s. Final extension at 72℃ for 5min.

Results: At 60℃ annealing, very bright band around 200 bp, 332 bp fragment is now visible but only very faint fragments where the control should be. At 50 and 55℃ annealing more larger bands appear but its too many and not clear which are the fragments we are looking for.

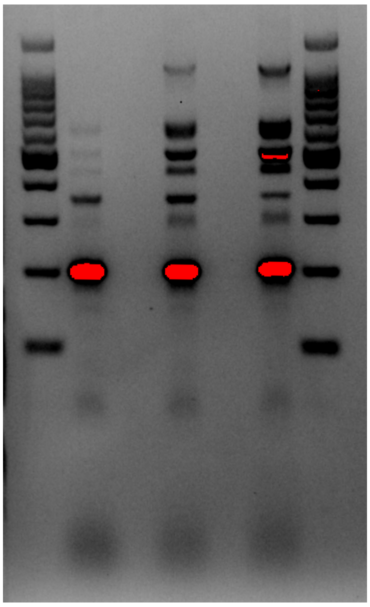

**Trail 3**

Going back to GoTaq but increasing $Mg^{2}+$ concentrations to 2.5/ 3.5/ 4.5 mM (compared to 1.5 mM in the original GoTaq buffer). PCR program of trial 1.

Results: Increasing $Mg^{2}+$ does make more fragments appear, but there are more than anticipated (something around 400bp). Control fragment(?) remains vague. One sample shows something that could be 332 bp.

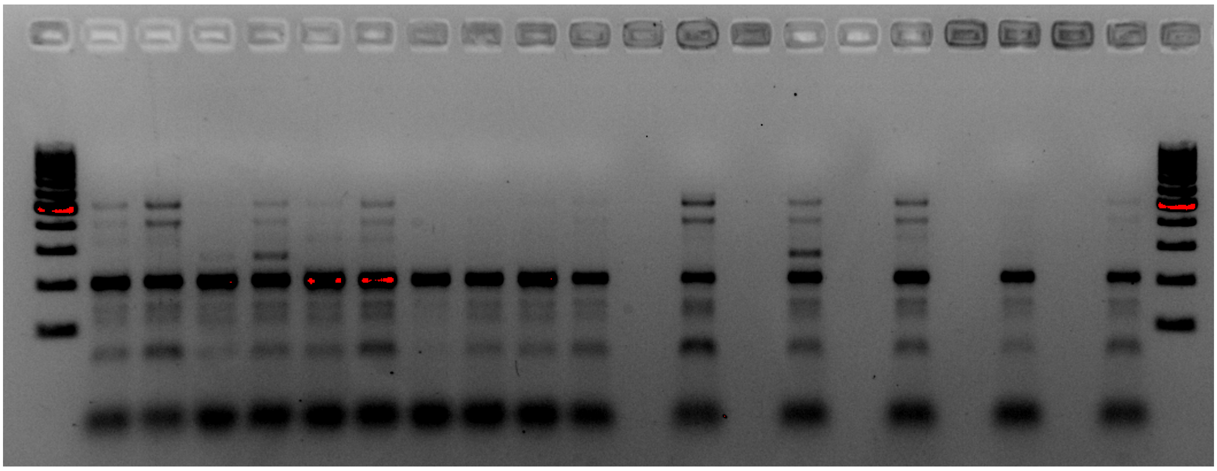

**Trial 4**

Try the exact same protocol as Panini et al. (2014) with Dreamtaq green PCR mastermix.

PCR program: 94C for 2min, 30 cycles of 94℃ for 30s, 48℃ for 30s and 72℃ for 30s. Final extension at 72℃ for 5min.

Results: No control fragment.

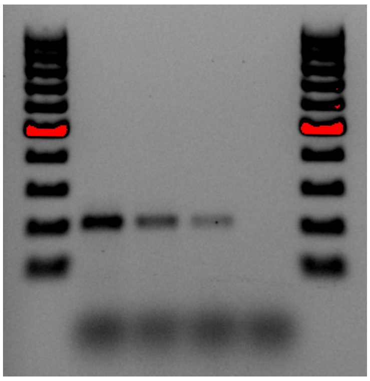

**Trial 5**

Only test primers MpNACR-R514 and MpNACR-F52 that should result in a 508 bp control fragment. Using Dreamtaq protocol (trial 4) but with 45s extension.

Results: Faint control band.

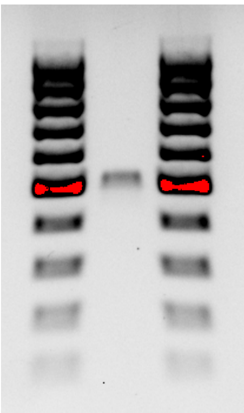

**Trial 6**

Use 1.5x of primers MpNACR-R514 and MpNACR-F52 compared to the other two primers. Test annealing at 61, 59, 57 and 55℃ and for 45s. Extension for 45s. Cycled for 32X. Test two DNA samples.

Results: At 57/59℃ annealing, both samples display the mutation fragment together with the control and wild-type fragments. Trial 6 shows potential.

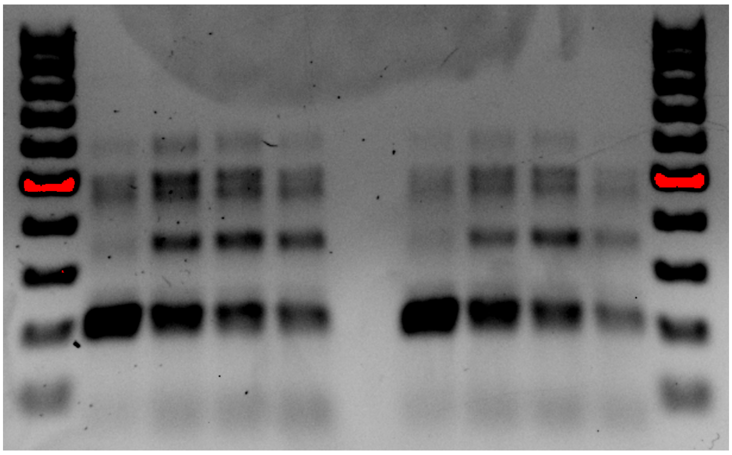

**Trial 7**

Methods of Trial 6 but with annealing at 58.5℃. Testing 13 DNA samples, all with different multilocus genotypes.

Results: All samples display all three fragments, which is not realistic because mutation R81T is not very common.

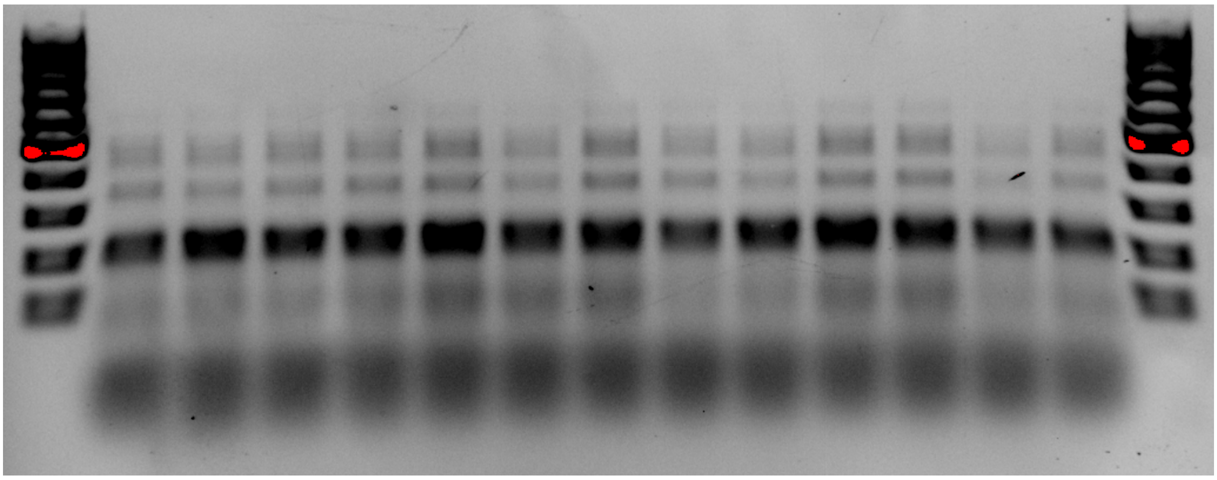

### Supplementary File S4: Number of aphids tested in the experiments

##### Pymetrozine

In the pymetrozine treatments, the number of aphids tested, control treatments excluded (Figure S4-1A), differed significantly between replicates (LM: $F(2, 120) = 7.87$, $p < .001$; Figure S4-1B), with an average of 37.2 for the first two replicates and 29.9 for the third. Additionally, the number of aphids tested also varied between aphid lines (LM: $F(8, 120) = 12.51$, $p < .001$; Figure S4-1C), with averages per aphid line ranging from 22.8 to 51.2. Lastly, the number of aphids tested also varied between the different concentrations of pymetrozine (LM: $F(4, 120) = 3.77$, $p = .006$; Figure S4-1D), with the averages ranging from 31.9 to 41.4.

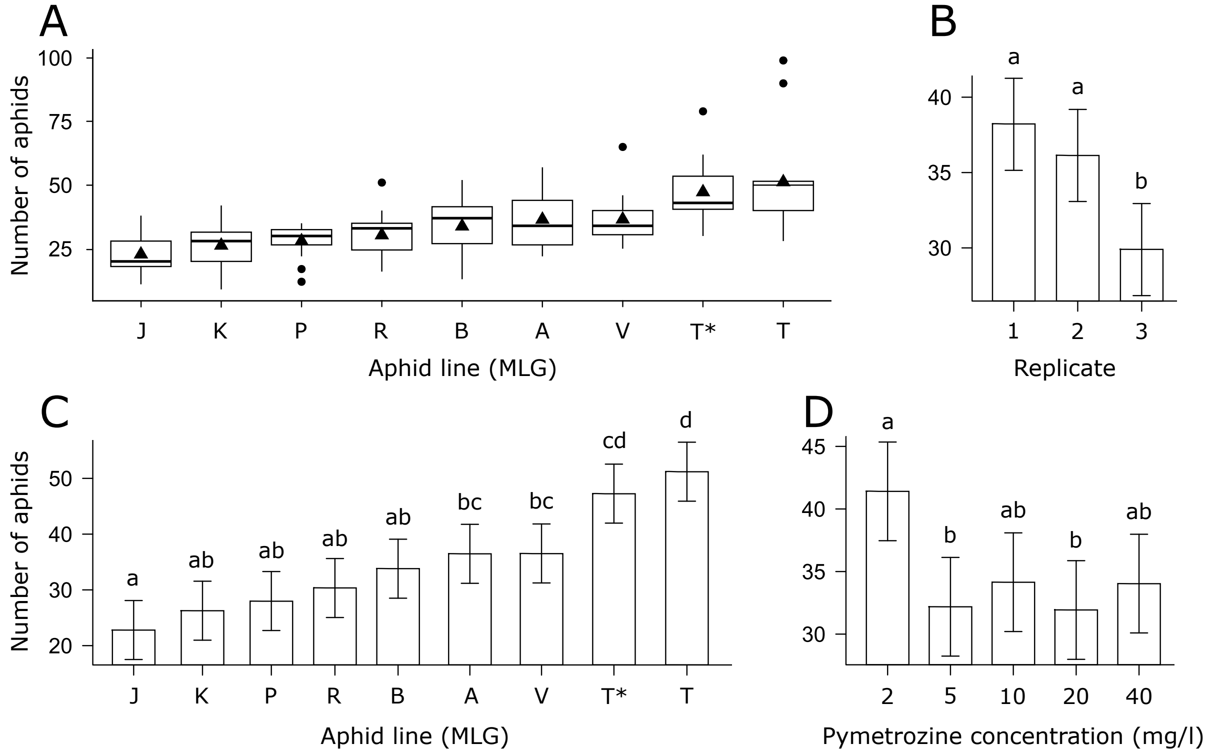


**Figure S4-1.** Pymetrozine experiment A) Boxplot displaying the number of aphids tested, ordered from lowest to highest mean. Triangles depict the mean and circles the outliers B) Bargraph displaying the effect of 'replicate' on the number of nymphs C) Bargraph displaying the effect of 'aphid line' on the number of nymphs, ordered from lowest to highest mean D) Bargraph displaying the effect of 'treatment (pymetrozine concentration)' on the number of nymphs. In panels B, C and D, the error bar represent the standard error and the letters depict in which treatments the number of aphids included deviated significantly ($p < .05$) from each other, based on Tukey's post hoc testing. The names of the aphid lineages are abbreviated according to the MLG of each line.

**Flonicamid experiment 1**

In flonicamid experiment 1, the number of aphids tested, control treatments excluded (Figure S4-2A), differed significantly between replicates (LM: $F(2, 120) = 34.31$, $p < .001$; Figure S4-2B), with an average of 30.8 aphids in the first two replicates and 19.7 in the third. Additionally, the number of aphids tested also differed between aphid lines (LM: $F(8, 120) = 9.45$, $p < .001$; Figure S4-2C), with averages per aphid line ranging from 20.7 to 39.4. However, the number of aphids tested did not differ between the different concentrations of flonicamid (LM: $F(4, 120) = 1.30$, $p = .276$; Figure S2D).

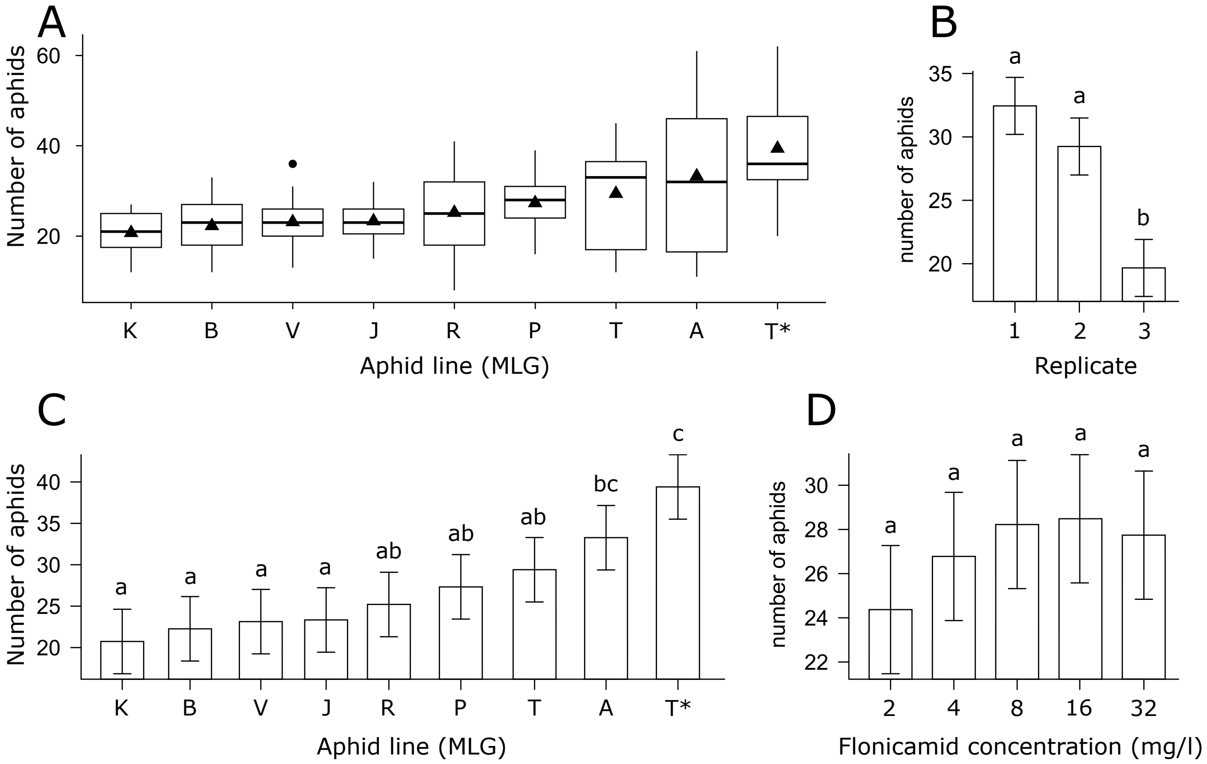


**Figure S4-2.** Flonicamid experiment 1. A) Boxplots displaying the number of aphids tested, divided by aphid line, ordered from lowest to highest mean. Triangles depict the mean and circles the outliers. B) Bargraph displaying the effect of 'replicate' on the number of nymphs C) Bargraph displaying the effect of 'aphid line' on the number of nymphs, ordered from lowest to highest mean D) Bargraph displaying the effect of 'treatment (flonicamid concentration)' on the number of nymphs. In panels B, C and D, the error bar represent the standard error and the letters depict in which treatments the number of aphids included deviated significantly ($p < .05$) from each other, based on Tukey's post hoc testing. The names of the aphid lineages are abbreviated according to the MLG of each line.

**Flonicamid experiment 2**

In flonicamid experiment 2, the number of aphids tested, control treatments excluded, did not differ between aphid lines (LM: $F(5, 27) = 2.46$, $p = .059$), concentrations of flonicamid (LM: $F(1, 27) = 0.48$, $p = .496$), nor replicates (LM: $F(2, 27) = 0.75$, $p = .481$)(Figure S4-3).

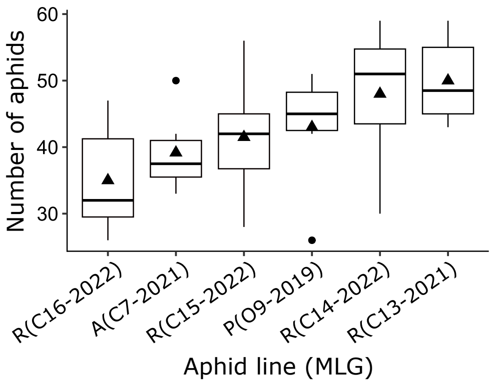


**Figure S4-3.** Boxplot displaying the number of aphids tested in flonicamid experiment 2, per aphid lineage, ordered from lowest to highest mean. Triangles depict the mean and circles the outliers.
